# Oro-manual assessment of food preservability in a wild rodent

**DOI:** 10.64898/2026.09.23.752726

**Authors:** Lucas Dumargne, Esther Tibi, Margaret Williams, Dumas Galvez, Camille Testard, Juan I Sanguinetti-Scheck

## Abstract

Animals encounter a variety of food items and must determine their value and preservability. How such assessments are made in natural contexts remains largely unknown. Here, we dissect how agoutis—a critical Neotropical seed disperser—decide whether to immediately consume or preserve food in small independent caches, i.e. scatter-hoard. Agoutis preferentially cached nuts over fresh fruits and underwent a sharp transition from eating to scatter-hoarding, a clear shift towards future oriented decision-making. Remarkably, they cached empty nuts with intact shells but consumed filled nuts with damaged shells, revealing that shell integrity, rather than immediate nutritional content, determines the decision to scatter-hoard. This decision depended on oro-manual food assessment, an ecological information-seeking behavior. Agoutis used their lips and incisors—a somatosensory fovea—to systematically navigate nut surfaces, accumulating decision-critical sensory information about shell integrity. Together, these findings provide a mechanistic account of preservability assessment in a wild rodent, revealing how information-seeking behaviors support long-term value optimization in natural environments.

## Introduction

To survive, animals must trade off current and future needs(*1*). This requires making so-called “intertemporal decisions” whereby they compare actions and their resulting rewards when they occur at different, remote points in time(*1*, *2*). Animals, including humans, are known to favor immediate smaller rewards over larger rewards in the distant future(*3–7*). However, some animals(*8*, *9*) may forgo immediate reward to prepare for potential future scarcity. These behaviors depend on accurately estimating a food item’s future preservability, yet the computations, behavioral algorithms, and mechanisms that support such assessments remain poorly understood(*10*, *11*). This gap persists because mechanistic research on foraging has been almost exclusively performed on lab rodents and under simplified, highly-controlled experimental setups with water or simplistic foods(*12–16*) such as pellets. While these paradigms provide experimental control, they largely eliminate the structural complexity of natural foods, the statistics of their occurrence in time and the rich sensory interactions required to evaluate them(*17–19*). In nature, foragers experience structured food items like fruits with peels, shells, linings, pulps and seeds. To optimize long-term value, animals must assess the affordances(*20*) (use properties) of encountered food items and their components such as their transportability, graspability, preservability and consumability. Extracting ecological information often requires active information-seeking actions(*20*) deployed to resolve environmental uncertainty, sometimes at the expense of immediate reward(*21*, *22*). In this study we investigate such processes *in natura* with high behavioral resolution and repeatability by harnessing the remarkable scatter-hoarding behavior of the agouti (*Dasyprocta punctata*) – a system that allows us to perform systematic task designs in wild animals(*17*) and to study the mechanics of food assessment behaviors.

Agoutis are large diurnal frugivorous rodents living in the rainforests of the Americas and have been extensively studied in behavioral ecology due to their critical role in maintaining tropical forest diversity(*23–25*). They are known for opening large seed pods in the rainforest and dispersing seeds for later use – an intertemporal decision termed « scatter-hoarding »(*9*, *26*, *27*). Scatter-hoarding consists in the active concealment of seeds, sometimes fruits, in different independent small caches in the soil to save food for future use(*9*). This behavior requires a trade-off between immediate and future needs, implying decision-making that integrates both short– and long-term utility. Previous work has begun to describe the agouti’s seed choice(*28–30*), cache distribution(*26*, *31*), inspection(*32*), retrieval(*32*) and contribution to forest maintenance(*23*, *33*, *34*). Evidence shows that scatter-hoarders use information about the nutritional quality of nuts when making cache decisions such as fat content, size or tannins contents(*29*, *30*, *35–37*) but the specific information seeking mechanisms that underlie this natural intertemporal decision-making (i.e. consume now or save for later) process remains largely unexplored.

Our study combined methods in behavioral ecology, computational ethology and modeling to investigate the mechanisms of preservability assessment in scatter-hoarding agoutis with unprecedented detail. We find that upon encountering food with the potential for storage, agoutis can transition into a scatter-hoarding behavioral state that dramatically changes their foraging ethogram to include preservability assessment, an information-seeking behavior. To decide which nuts to store, agoutis perform a coordinated oro-manual navigation of the nut’s outer shell to detect breaches using their lips and teeth, their somatosensory fovea(*38*). Field experiments confirmed that this assessment behavior leads agoutis to cache nuts with higher preservability and lower pilferage risk. Their assessment behavior conforms to nuts’ shape, optimizes search efficiency, and parallels active sensing in vision(*39*, *40*). Overall, our work reveals how haptic information-seeking supports preservability assessment and intertemporal decision-making in a wild scatter-hoarding mammal.

## Results

### The agouti’s scatter-hoarding behavior undergoes a sharp state transition

Central American Agoutis (*Dasyprocta punctata*) (***Fig. 1A***) are found in Central and South America (***Fig. 1B***). We studied a wild population of agoutis living in Gamboa, Panama (***Fig. 1B***), a small town connected to the Soberania National Park. Even under human presence, agoutis are avid scatter-hoarders, a complex behavior composed of a long behavioral sequence: animals first locate a food item, they then transport it far from its source, dig a hole in the ground, place the item in the ground, and finally cover the hole with soil, often placing a leaf on top of the cache (***Fig. 1C, Supplementary video 1***). We recorded agoutis’ scatter-hoarding behavior of provided fruits using camera traps set up across Gamboa and observed 251 agouti-fruit interactions (***Fig. 1D-E, S1A***). Agoutis scatter-hoarded fruits provided at different rates: large fleshy fruits like bananas were rarely scatter-hoarded while small and dry fruits like grapes or peanuts were more frequently scatter-hoarded (peanut-apple: p-value < 0.001, peanut-banana: p-value < 0.001, peanut-grape: p-value < 0.01, Two-sample Z-test Bonferroni-corrected; ***Fig. 1F***). Overall, agoutis preferred scatter-hoarding shelled fruits and nuts, which have higher preservability(*41*). Based on this finding we focus on peanuts for the rest of this study.

**Figure 1:**
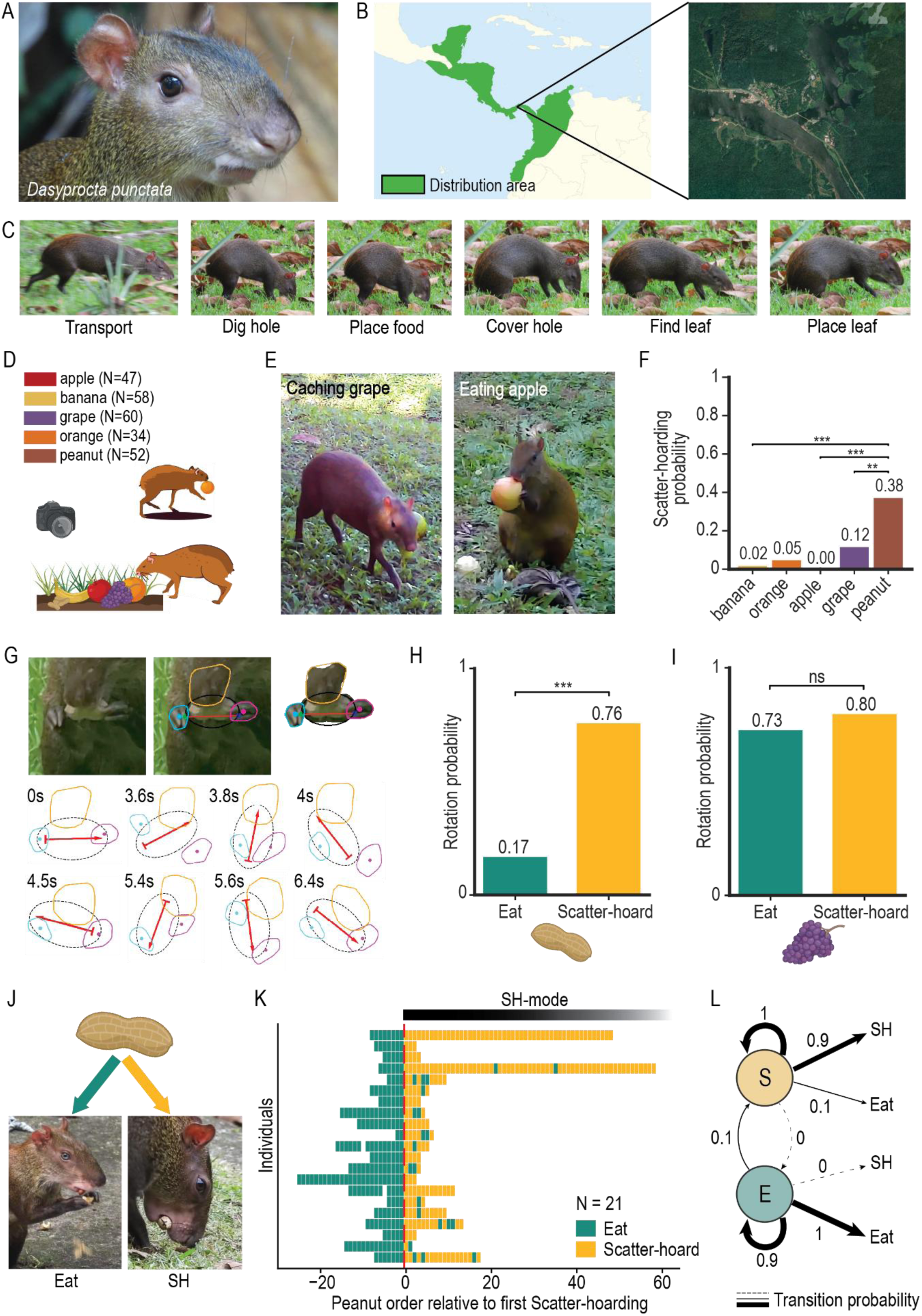
Agouti scatter-hoarding decision-making. **A.** Central American Agouti (*Dasyprocta punctata*). **B.** Field location in Gamboa, Panama. **C.** Agouti scatter-hoarding temporal ethogram: food transport, hole digging, food placing, hole covering, leaf finding and placing. **D.** 251 agouti-fruit interactions were recorded (47 apples, 58 bananas, 60 grapes, 34 oranges, and 52 peanuts). **E.** Agouti caching a grape (left) and eating an apple (right). **F.** Scatter-hoarding probability for all fruits provided (peanut-apple: p-value < 0.001, peanut-banana: p-value < 0.001, peanut-grape: p-value < 0.01, Two-sample Z-test, Bonferroni-corrected). **G.** Tracking of extensive peanut rotation (peanut: black outline, long axis: red arrow, mouth: orange outline, right hand: blue outline, left hand: pink outline), cf. ***Supplementary video 2***. **H.** Rotation probability given the decision to eat (teal) and scatter-hoard (yellow) for 56 peanuts (P(Rotation|Scatter-hoard) = 0.76, P(Rotation|Eat) = 0.17; p-value < 0.001, Two-sample Z-test). **I.** Same as **H.** for 84 grapes (P(Rotation|Scatter-hoard) = 0.80, P(Rotation|Eat) = 0.73; p-value > 0.05, Two-sample Z-test). **J.** Agoutis were given a series of peanuts (one trial) and chose to eat or scatter-hoard them. **K.** 21 trials (rows, rectangles represent peanuts, teal if eaten, yellow if scatter-hoarded) aligned on the first scatter-hoarding event. **L.** Transition and emission probabilities of the best HMM fitted to empirical data. In all the figure, bars correspond to the mean probability of behavior per fruit, *: p-value < 0.05, **: p-value < 0.01, ***: p-value < 0.001, ns: not significant.

Observations from camera traps revealed that agoutis sometimes extensively rotated peanuts before making a foraging decision (***Fig. 1G, Supplementary video 2***), raising the possibility that rotation serves as an ecological information-seeking behavior related to scatter-hoarding. We therefore asked whether this behavior was associated with the eventual outcome of the manipulation. Strikingly, peanuts that were eventually scatter-hoarded were significantly more likely to be rotated than peanuts that were consumed (N = 56 peanuts, p-value < 0.001, Two-sample Z-test; ***Fig. 1H***). In contrast, this was not the case for similarly sized grapes, where no significant difference was found (N = 84 grapes, p-value > 0.05, Two-sample Z-test; ***Fig. 1I***), indicating that the association between rotation and scatter-hoarding is not a general feature of handling small food items but is specific to scatter-hoarding of peanuts. Because of the opportunistic nature of the camera-trap data, we could not determine whether rotation promoted scatter-hoarding or whether the decision to scatter-hoard increased the likelihood of rotation.

To distinguish these possibilities, we experimentally induced scatter-hoarding in identified individuals (using visual markings; ***Fig. S1C***) and then provided them with whole, intact peanuts one after the other, allowing them to choose whether to eat or scatter-hoard (***Fig. 1J***). Early in the experiment all individual agoutis (N = 21) exhibited a tendency to consume the peanuts but subsequently underwent a sharp transition from this “eating” mode to “scatter-hoarding” (henceforth “SH”). Following the first SH epoch, scatter-hoarding events sharply increased in frequency and became the predominant behavioral state (***Fig. 1K***), hereafter referred to as “*SH-mode*”. This state persisted for the rest of the experiment. The abruptness and persistence of this behavioral shift suggested that scatter-hoarding might reflect a discrete internal state rather than a gradual change in behavioral preference. This was confirmed by a modeling approach, showing that a two-states Hidden Markov Model (HMM) with an Eating-Mode state and a SH-mode state, both emitting ‘Eat’ and ‘SH’ behaviors, showed the best fit among other more progressive models (***Fig. 1L, S2A-B***, see methods). This best fit HMM (***Fig. S2C***) consisted of a very absorbent, persistent SH-mode state that never reverts to Eating-mode within the timeframe of our experiments (transition probability: S → E = 0; ***Fig. 1L, S2D***). Furthermore, exploration of the HMM parameter space revealed that the best fit was found for very low values of q (probability of SH while in Eat-mode) and high values of r (probability of SH while in SH-mode) (***Fig. S2E***), showing that this process relies on the stable establishment of a SH-mode rather than on noisy emission probabilities of behaviors (***Fig. S2E-F***). Overall, our results show that agoutis undergo a state transition from eating to a persistent (absorbent) SH-mode.

### Shell integrity assessment as a computation of food preservability

Next, we investigated the role of peanut rotations in scatter-hording decisions of wild agoutis. To do so, in addition to the outcome of the Eat vs SH decision, we scored whether a rotation behavior was performed for each peanut (***Fig. 2A***). We found a drastic increase of rotations after the switch into SH-mode (N = 21 individuals, pre-switch = 0.05, post-switch = 0.48, p-value < 0.001, Two-sample Z-test) (***Fig. 2B***). After the switch however, the rotation probability is the same for eaten and cached peanuts (eaten = 0.41, scatter-hoarded = 0.49, p-value > 0.05; ***Fig. S2F***), indicating that the rotation increase is related to the state of SH-mode, rather than the behavioral outcome of the decision. A bivariate version of the aforementioned two-state HMM, where Eat– and SH-modes emit rotation behaviors in addition to the outcomes (***Fig. S2G***), explains the empirical data well (***Fig. S2K-L***) for both the decisions (***Fig. S2J***) and the rotations (***Fig. S2H***), while resulting in the same sharp state transition (***Fig. S2M***). This data is consistent with rotations as a state-dependent assessment behavior related to the act of scatter-hoarding for peanuts. Next, we asked whether assessment could be an ecological information-seeking behavior for agoutis to evaluate the internal nutritional value of the peanuts before making their decision.

**Figure 2:**
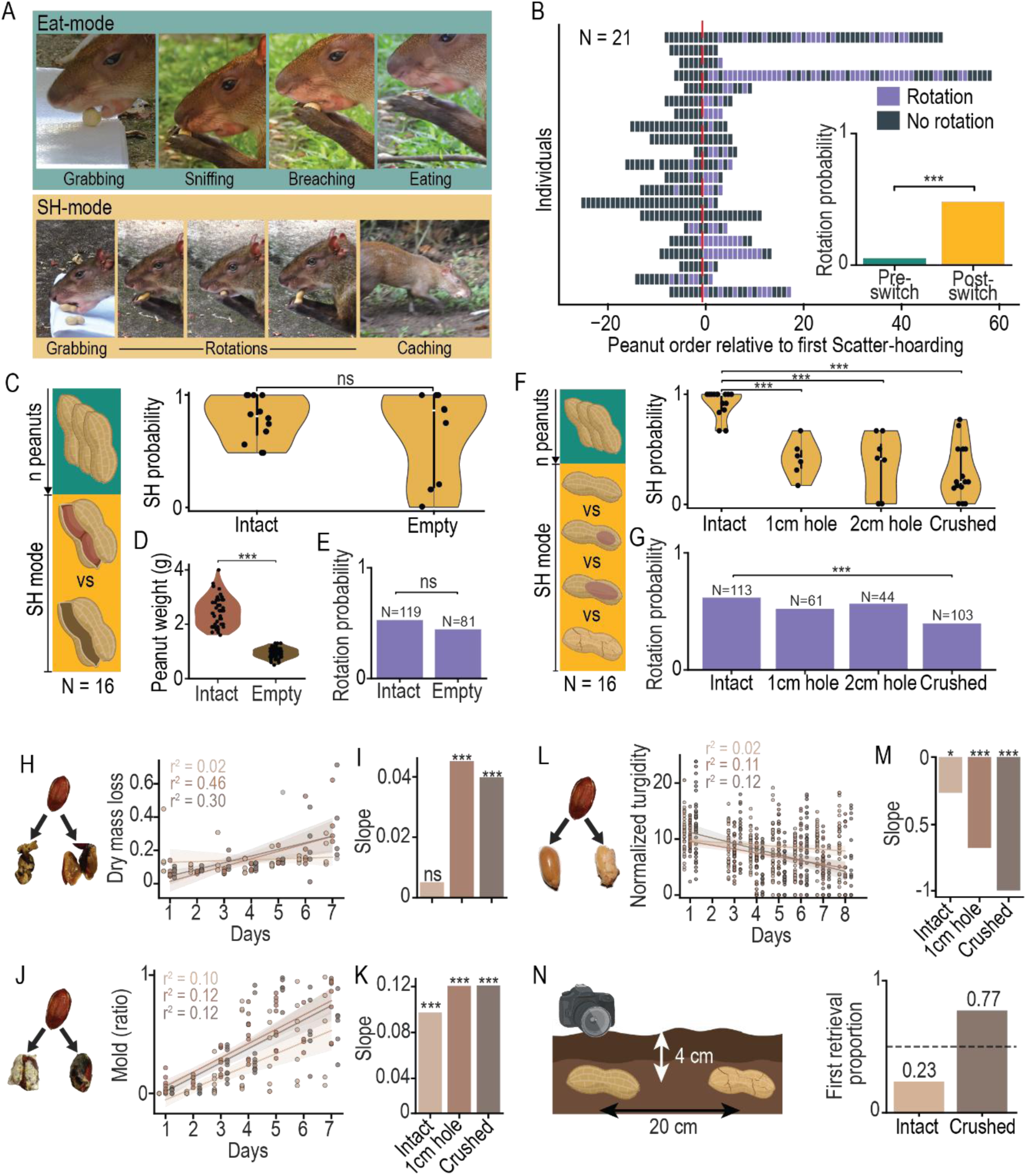
Agoutis assess peanut preservability. **A.** Pre-eating and pre-caching ethograms. **B.** Same as **1K.** but colored by whether the peanut was rotated (purple) or not (grey) and overall probability of rotation before (teal) and after (yellow) switch to SH-mode (p-value < 0.001, Two-sample Z-test). **C.** 16 agoutis in SH-mode are given intact peanuts or peanuts that were emptied and reshelled. Distributions of SH probabilities across individuals for each peanut condition (p-value = 0.83, two-sided Mann-Whitney U test). **D.** Weight distributions for intact and empty peanuts (Intact: 2.5, Empty: 0.9, p-value < 0.001, two-sided Mann-Whitney U test). **E.** Rotation probability for intact and empty conditions (p-value = 0.25, two-sided Fisher’s exact test). **F.** 16 agoutis in SH-mode are given peanuts whose shells are intact, present a hole of 1 cm, 2 cm, or are crushed. Distributions of SH probabilities across individuals for each condition (significant pairs: intact-1cm hole p-value < 0.001, intact-2cm hole p-value < 0.001, intact-crushed p-value < 0.001, pairwise Fisher’s exact test Holm-corrected after omnibus *χ*^2^ – p-value < 0.001). **G.** Rotation probability for each peanut condition (p-value < 0.001, pairwise Fisher’s exact test Holm-corrected after omnibus *χ*^2^ – p-value < 0.05). **H.** Perished peanuts after 4 days. Linear regression of dry mass loss for each retrieval time and condition (intact: r² = 0.02, 1cm hole: r² = 0.46, crushed: r² = 0.30). **I.** Slopes of the linear regression for each condition (intact: slope = 0.005, p-value > 0.05, 1cm hole: slope = 0.045, p-value < 0.001, crushed: slope = 0.040, p-value < 0.001). **J.** Molded peanuts after 4 days. Same as **H.** for the proportion of pixels containing mold on the seed (intact: r² = 0.10, 1cm hole: r² = 0.12, crushed: r² = 0.12). **K.** Same as **I.** for the regressions on proportion of molded pixels (intact: slope = 0.09, p-value < 0.001, 1cm hole: slope = 0.12, p-value < 0.001, crushed: slope = 0.12, p-value < 0.001). **L.** Differentially turgid peanuts after 4 days. Same as **H.** and **J.** for seed turgidity (intact: r² = 0.02, 1cm hole: r² = 0.11, crushed: r² = 0.12). **M.** Same as **I.** and **K.** for the regressions on normalized turgidity (intact: slope = –0.27, p-value < 0.05, 1cm hole: slope = –0.68, p-value < 0.001, crushed: slope = –0.99, p-value < 0.001). **N.** Differential perishability experiment. Proportion of peanuts retrieved first in a peanut pair for each condition (intact: probability = 23%, crushed: probability = 77%, dashed line: chance level). In all the figure, N: number of individuals in experiment (black dot: within-individual average, violin: between-individual distribution), purple bars correspond to the mean probability of rotation (N: number of peanuts, brown dot: peanut, line: best fitted line of a linear regression, shaded envelope: CI 95, r^2^: regression’s coefficient of determination). *: p-value < 0.05, **: p-value < 0.01, ***: p-value < 0.001, no bracket: not significant.

To test this hypothesis, we quantified the foraging decisions of agoutis already in SH-mode when presented with either intact peanuts or empty re-shelled peanuts (***Fig. 2C***). Surprisingly, agoutis continued to scatter-hoard empty peanuts with probabilities comparable to intact peanuts (N = 16 individuals, p-value = 0.83, two-sided Mann-Whitney U test; ***Fig. 2C***), despite intact peanuts being more than twice as heavy as empty peanuts (p-value < 0.001, two-sided Mann-Whitney U test; ***Fig. 2D***). We observed no significant difference in rotation probability between the two conditions (p-value = 0.25, two-sided Fisher’s exact test; ***Fig. 2E***) implying consistent assessment. These findings suggest that, once in SH-mode, agoutis did not rely primarily on internal value.

We hypothesized that, instead, agoutis used rotations to evaluate shell integrity, which could affect the preservability affordances of shelled food items. To test this hypothesis, we presented agoutis in SH-mode with peanuts spanning a gradient of shell integrity: intact shells, shells containing a 1 cm breach, shells containing a 2 cm breach, and crushed shells with extensive cracks (***Fig. 2F, S3A)***. Even while in SH-mode agoutis were selectively less likely to scatter-hoard peanuts with lower shell integrity (N = 16 individuals, significant pairs: intact-1cm hole p-value < 0.001, intact-2cm hole p-value < 0.001, intact-crushed p-value < 0.001, pairwise Fisher’s exact test Holm-corrected after omnibus *χ*^2^-p-value < 0.001; ***Fig. 2F***). The probability of rotating peanuts remained unchanged across shell conditions, except for crushed shells, which were rapidly assessed, breached and consumed (significant pair: intact-crushed p-value < 0.001, pairwise Fisher’s exact test Holm-corrected; ***Fig 2G***). We found similar results for a naturally occurring palm seed (*Attalea butyracea)* confirming our results generalize across shelled food items (***Fig. S3B-C***). Our results reveal that shell integrity, and not assessment effort, is the critical factor in SH decision-making.

We next asked whether the preference for intact peanuts for scatter-hoarding was ecologically beneficial. We hypothesized that shell integrity serves as a cue for preservability and future value, with breached peanuts being less likely to retain value after being cached. We tested two major risks faced by stored resources in the wild: deterioration and pilferage. To quantify deterioration risk, we buried peanuts with intact, breached, or crushed shells and monitored their condition for up to seven days. Peanuts with compromised shells deteriorated more rapidly than intact peanuts. They lost dry mass faster (intact: r² = 0.02, slope = 0.005, 1cm hole: r² = 0.46, slope = 0.045, crushed: r² = 0.30, slope = 0.040; ***Fig. 2H-I***), accumulated mold more rapidly (intact: r² = 0.10, slope = 0.09, 1cm hole: r² = 0.12, slope = 0.12, crushed: r² = 0.12, slope = 0.12; ***Fig. 2J-K***), and exhibited a stronger decline in turgidity (intact: r² = 0.02, slope = –0.27, 1cm hole: r² = 0.11, slope = –0.68, crushed: r² = 0.12, slope = –0.99; ***Fig. 2L-M***) than intact counterparts after three days (N = 30 peanut per day per condition per experiment). This demonstrates that breached peanuts have shorter preservability and therefore lower long-term value. To measure pilferage, we manually buried pairs of intact and crushed peanuts mimicking natural agouti caches (***Fig. 2N***). Crushed peanuts were retrieved before intact peanuts in 77% of trials (N = 26 pairs; ***Fig. 2N***), demonstrating that shell damage substantially increases vulnerability to pilferage, probably mediated by a more conspicuous smell. Overall, our results indicate that shell integrity reliably predicts the future fate of cached peanuts in the wild. Thus, the cues agoutis use during shell assessment accurately indicate long-term value of food items. Therefore, pre-scatter-hoarding assessment behavior appears to be a targeted information-seeking behavior allowing animals to assess preservability.

### Oro-manual algorithm for preservability assessment in agoutis outperforms canonical search algorithms

To uncover the algorithms that agoutis use to obtain ecological information about preservability, we manually scored 282 videos creating a comprehensive ethogram of their assessment (N = 15; ***Fig 3A-B***). We observed the following behaviors: rotations (using mouth or forepaws), jaw clamps (tightly biting the peanut with their teeth without breaching), licking, sniffing, and mouth-to-hole contact (***Fig. 3A-B, Supplementary video 3***). Analysis of transitions between these behaviors revealed a highly structured assessment sequence dominated by alternating bouts of rotation and jaw clamps (***Fig. 3A, S4A***). Notably, rotations were highly likely to be followed by jaw clamps (***Fig. 3C***), revealing a tight coordination between forelimb manipulation and oral probing during assessment.

**Figure 3:**
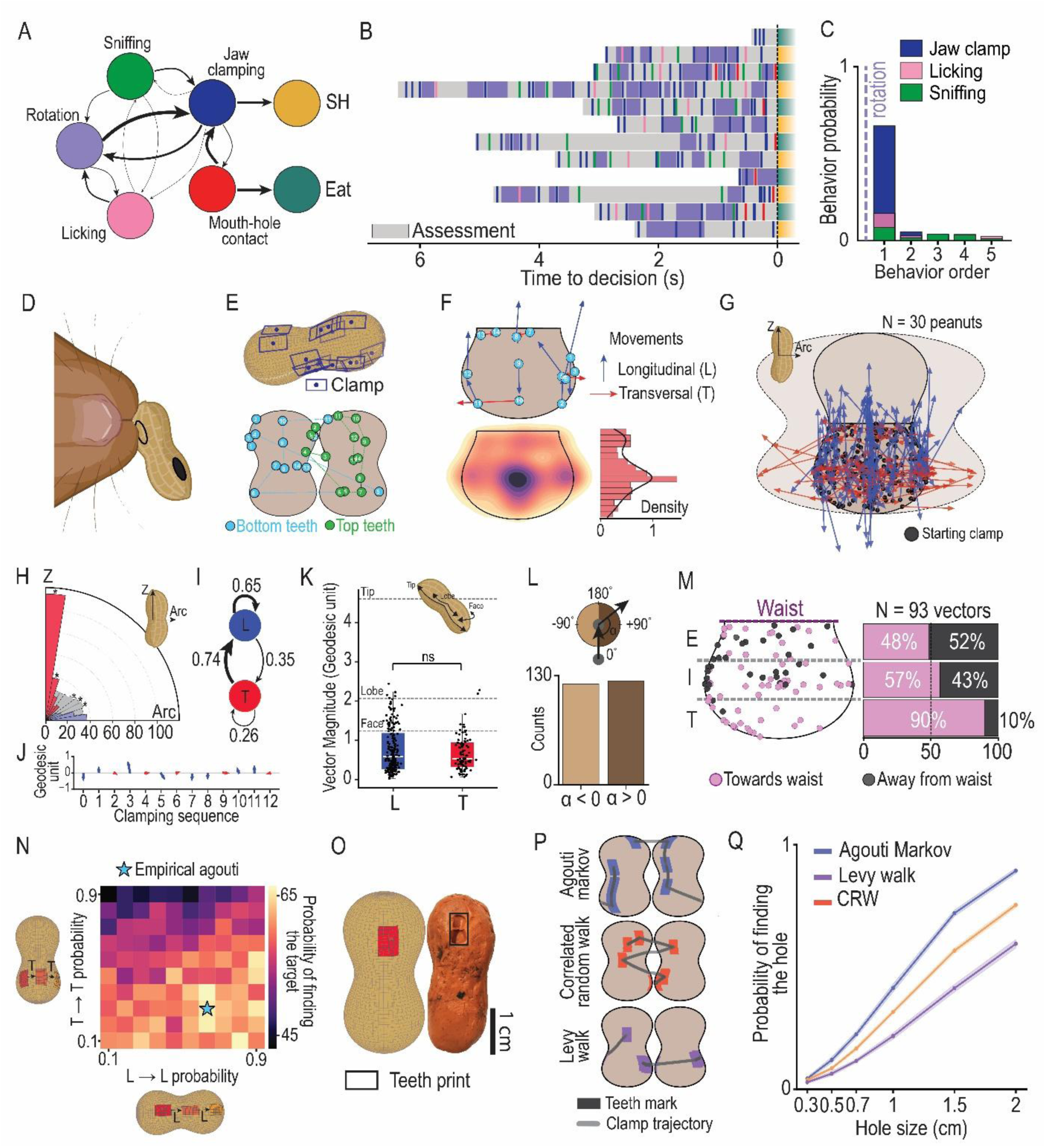
Agoutis strategically navigate shells during assessment. **A.** Assessment ethogram (arrow: transition probability). **B.** Scoring of assessment behaviors for example peanuts (row: peanut, colors after time 0s: decision outcome, teal: eat, yellow: SH). **C.** Probability of jaw clamp (dark blue), sniffing (green), and licking (pink) after a rotation. **D.** Peanut’s visual landmarks. **E.** Example of a jaw clamping sequence on a real-scale 3D peanut model and inferred 2D clamping map for bottom (light blue) and top (light green) teeth. Numbers correspond to the order of the clamp in the sequence. **F.** Minimal map of clamp-to-clamp vectors for an example sequence (top, dark blue: longitudinal vectors (L), red: transversal vectors (T)) and density of clamps across all peanuts (bottom, N = 30). **G.** Same as **F. top** with data compiled over 30 peanuts (dark blue: longitudinal vectors (L), red: transversal vectors (T)). **H.** Distribution of clamping vector orientations across 10 angle bins compared to null distribution (*χ*2 test: p-value < 0.001, post-hoc bin analysis: [81°-90°]: p-value < 0.001, [72°-81°]: p-value < 0.05, [36°-45°]: p-value < 0.001, [27°-36°]: p-value < 0.01, [18°-27°]: p-value < 0.05, others are not significant). **I.** Transition probabilities graph for L (blue) and T (red) vectors. **J.** Timeline of clamping vectors for an example peanut. **K.** Magnitudes of L and T vectors in across all sampling sequences relative to tip-to-tip (Tip), face-to-face (Face), lobe-to-lobe (Lobe) distances (N = 305 vectors, geodesic distance, p-value > 0.05, two-sided Mann-Whitney U test). Black scatters are individual vectors. **L.** Computation method for vector-to-vector angles and count off left– and right-wards turns (N = 263 vector pairs, stats). **M.** Location distribution of longitudinal vectors (N = 93) oriented towards (pink) or away from (grey) the waist (purple dashed line) in three regions along the longitudinal axis of a peanut (E: Equatorial, I: Intermediate, T: Tip) and share of the vectors in these three regions. **N.** Probability of finding a 1 cm diameter target on the ovoid shape across a sweep of T → T and L → L transition probabilities (blue star: Empirical agouti data). **O.** Bottom teeth print on clay peanut (back rectangle) and simulated teeth clamp on the 3D model of peanut (red vertices). **P.** Examples of simulated walks (data-derived Markov: dark blue, Levy Walk: purple, Correlated Random Walk – CRW: dark orange) sampling the surface of the peanut shell using clamps (colored rectangles). **Q.** Probability of clamp-hole contact across 10,000 simulations for each type of walk described in **P.**.

To investigate how agoutis explore the peanut’s shell, we used artificial visual landmarks on the shell (***Fig. 3D, S4B, Supplementary video 4***) in combination with high zoom cameras to reconstruct the precise locations of successive jaw clamps events and map them on a 2D and a 3D model of a peanut (***Fig. 3E***). This allowed us to extract the movement vectors of the position of the mouth from one clamp to the next. Across 30 intact peanuts, we first observed that agoutis preferentially clamp the center of a peanut chamber (***Fig. 3F***). Second, we found that agoutis navigate the shell surfaces non-randomly, showing a strong preference for longitudinal and transversal movements (***Fig. 3F-H***) while avoiding diagonal trajectories (*χ*^2^ test: p-value < 0.001, post-hoc bin analysis: [81°-90°]: p-value < 0.001, [72°-81°]: p-value < 0.05, [36°-45°]: p-value < 0.001, [27°-36°]: p-value < 0.01, [18°-27°]: p-value < 0.05, others are not significant; ***Fig. 3H***). Third, the sequence of movements was itself structured: longitudinal movements preferentially followed one another, whereas the transversal movement most often transitioned into longitudinal ones (T → L: p = 0.74, L → L: p = 0.65, L → T: p = 0.35, T → T: p = 0.26; ***Fig. 3I***). Thus, the average probing sequence consists of repetitions of two longitudinal movements followed by a transversal movement (***Fig. 3J,*** for an example peanut), perhaps mirroring the peanut’s elongated geometry. To test whether agoutis adjust their probing movements to the peanut’s three-dimensional shape, we quantified the amplitudes of longitudinal and transverse movements and asked whether they differ according to the peanut’s anisotropy (i.e. geometric asymmetry). While longitudinal and transversal movements were not significantly different in their average magnitudes, the variance of longitudinal movements was higher (mean comparison: p-value > 0.05, Mann-Whitney U Test, L variance = 0.56, T variance = 0.43, variance comparison < 0.05, Levene’s test; ***Fig. 3K***). This greater variability is consistent with agoutis making a wider range of movement lengths along the peanut’s longer axis while maintaining more stereotyped movements across its shorter axis. We next asked whether agoutis exhibit stereotyped turning behavior during probing. To do so, we measured the angle between successive movement vectors and found no bias toward leftward or rightward turns (***Fig. 3L***). Instead, turning behavior depended on position on the peanut’s longitudinal axis: once reaching one of the peanut’s tips, agoutis were significantly more likely to turn toward the waist of the shell (***Fig. 3M***). This indicates that the direction of the next movement is guided by the animal’s location on the peanut rather than by an intrinsic turning preference or sequence. Together, these findings indicate that shell assessment is an active, spatially organized process in which agoutis continuously adjust their probing movements to the geometry of the peanut, rather than randomly sampling its surface.

Next, we asked whether seed navigation asymmetries would improve integrity assessment. To answer this, we simulated a range of navigation strategies that deviate from our data-driven Agouti Markov walk in their navigational asymmetries. For fixed average clamp vector magnitudes (longitudinal = 0.8 geodesic units, transversal = 0.8 geodesic units) we quantified the probability of finding a 1 cm diameter target on an ovoid shape across a sweep of T → T (transversal then transversal) and L → L (longitudinal then longitudinal) transition probabilities (***Fig. 3N***). The ovoid shape is a simplistic model of a real peanut designed to recapitulate its geometric asymmetry. We found that the navigational asymmetry empirically measured during assessment in agoutis (L → L: p = 0.65, R →T: p = 0.26) sits very close to the optimal strategy for finding the target on an ovoid (L → L: p = 0.65, T → T: p = 0.30; ***Fig. 3N***). Furthermore, our space exploration showed that strategies with higher L → L transition probabilities and lower T → T transition probabilities tend to perform better at finding a 1 cm target on an ovoid object. We also looked at how three navigational strategies with different asymmetries (agouti-derived, no asymmetry, agouti-inversed) perform in finding a target on a ovoids with peanut anisotropy and inverse anisotropy (***Fig. S4C-E***, see methods). In both ovoids, the best performing navigational strategy is the one whose most frequent movements align with the longest axis of the object (***Fig. S4D-E***). Overall, these results suggest that navigating an object in an asymmetric way that adapts to its anisotropy contributes to an optimal search.

Having established the importance of geometry-aware navigation, we next asked how the search algorithm employed by agoutis compares with canonical search strategies. Is geometry-aware navigation a strategy that makes assessment more efficient? To do so, we compared our data-driven Markov walk, augmented with the positional turning bias observed in ***Fig. 3M***, against two canonical search algorithms: a Correlated random Walk(*42*) (CRW) and a Levy walk(*43*) (***Fig. 3P***, see methods). Both these navigational strategies are proposed as optimal ways to explore a 2D manifold with unknown resource distribution what in the foraging literature(*42–44*). For each walk type, we simulated 10,000 sampling sequences and computed the probability of clamps (***Fig. 3O***) touching a hole of different diameters in a realistic 3D model of the peanut (***Fig. 3Q,*** see methods). We observed that the data-driven Markov walk outperforms the other navigational strategies in finding a hole across different sizes when looking for holes on the surface of a simulated peanut. We also compared the agouti strategy to a walk with memory (Self-Avoiding Walk – SAW, ***Fig. S4F-G***). Though the agouti Markov outperforms the SAW as well, SAW’s high performance suggests further that an organized navigational strategy improved finding a peanut hole. A binomial logistic regression with Markov walk as baseline showed the following odds ratios of success: Levy walk = 0.61, CRW = 0.92, SAW = 0.93. These findings indicate that the agouti’s geometry-aware navigational strategy, relying on a combination of directional asymmetry and position-dependent turning, allows them to outperform classic search algorithms at detecting holes on peanut shells.

### Mouth to hole contact underlies the decision to scatter-hoard

Having established how agoutis efficiently search peanut shells for breaches, we next asked how this information is translated into a decision to scatter-hoard or consume a peanut. Because holes are the primary cue predicting shell integrity, we hypothesized that direct contact between the mouth and a hole provides the sensory information that triggers this decision. To test this hypothesis, we presented agoutis with peanuts containing holes of different diameters spanning the range examined in our simulations (N = 11 individuals; ***Fig. 4A***). During shell exploration, repeated jaw clamping along the peanut surface enables extensive tactile probing with the teeth and lips, with some jaw clamps resulting in direct mouth-to-hole contact (***Fig. 3A, 4B, S4A, Supplementary video 5***). To quantify these events, two independent observers, annotated every instance in which the teeth or lips contacted a hole. Overall, 12% of scored jaw clamps resulted in observed mouth-hole contacts (***Fig. 3B, S4A***) and contact probabilities significantly increase as the holes increase in size (N = 11 individuals, r^2^ = 0.96, p-value < 0.01, linear regression; ***Fig. 4C***). Consistent with our understanding of agouti search, rotations increased the probability of mouth-hole contacts (p-value < 0.05, two-proportion Z-test Benjamini-Hochberg-corrected; ***Fig. 4D***). Detection of a hole using the mouth generally happened right before the start of eating (***Fig. 4E, S5C***) and resulted in a drastic increase in the probability of eating (p-value < 0.001, two-sided Fisher’s exact test, ***Fig. 4G***). Overall, we found that the probability of scatter-hoarding a peanut significantly linearly decreased with the size of the holes (N = 11 individuals, r^2^ = 0.34, p-value < 0.001, ***Fig. 4F***) demonstrating a highly sensitive perceptual process. These results suggest that detecting a hole is the main driver of the decision to eat. To further test this hypothesis, we applied a Generalized Linear Mixed Model (GLMM) to the behavioral data and outcomes. It revealed that mouth-hole contact was the only significant predictor of eating outcome among all assessment behaviors (coefficient = –0.278, p-value < 0.001, accounting for individual variability and varying hole diameters; ***Fig. 4H***). Together, these findings indicate that agoutis use coordinated rotation and jaw-probing behaviors to detect breaches in peanut shells, with mouth-hole contact serving as the critical sensory event that drives the decision to eat rather than cache.

**Figure 4:**
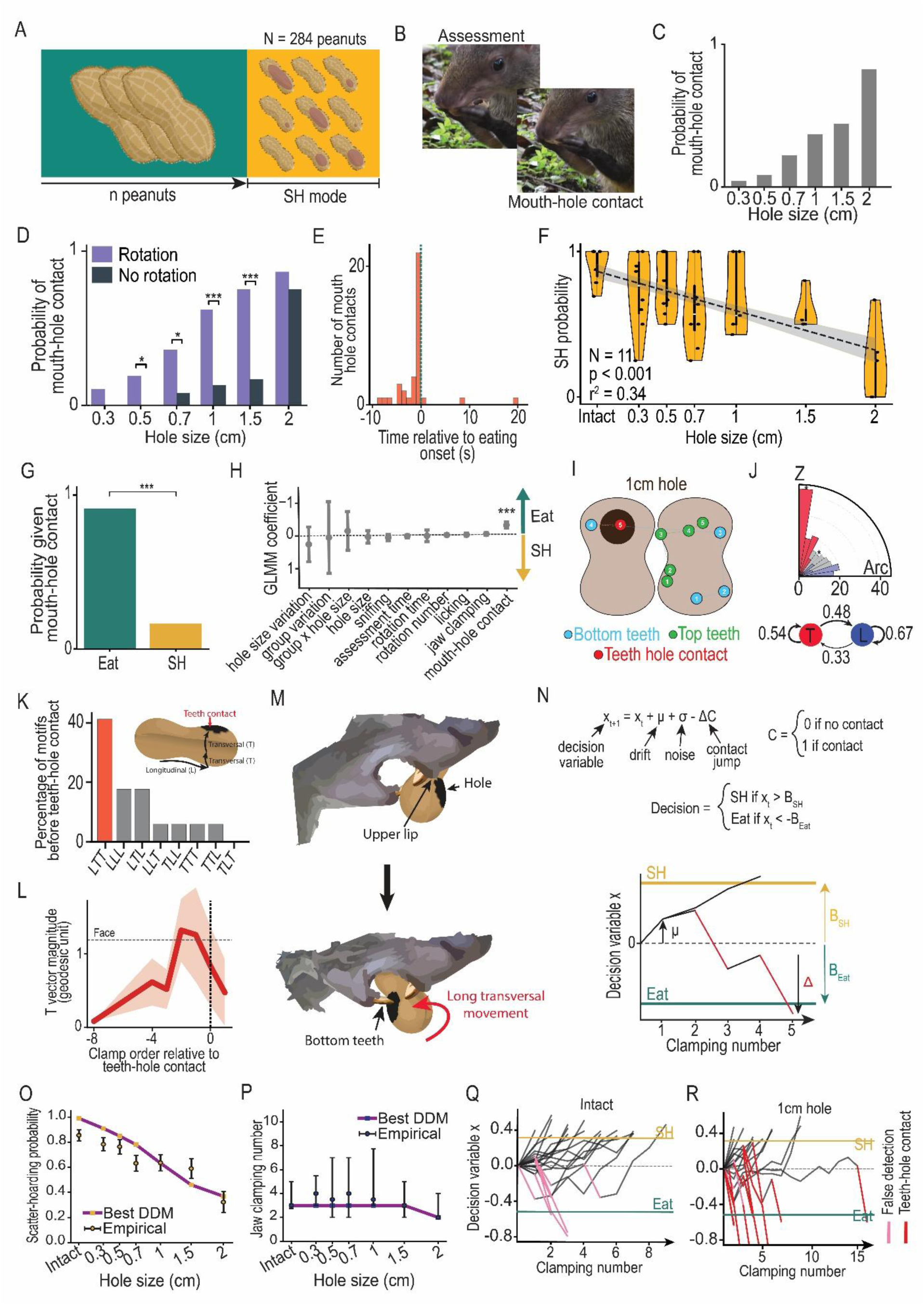
Decision is determined by mouth-hole contact. **A.** Experimental procedure: 11 individuals were given peanuts intact and with holes in their shells (diameters: 0.3, 0.5, 0.7, 1, 1.5, 2, 2.5 cm). **B.** Example of mouth-hole contact. **C.** Probability of mouth-hole contact across shell hole sizes (linear regression: r^2^ = 0.96, p-value < 0.01). **D.** Probability of mouth-hole contact across shell hole sizes given that the peanut was rotated (purple) or not (grey) (0.3 cm: p-value > 0.05, 0.5cm: p-value < 0.05, 0.7cm: p-value < 0.05, 1cm: p-value < 0.001, 1.5cm: p-value < 0.001, 2cm: p-value > 0.05, two-proportion Z-test Benjamini-Hochberg-corrected). **E.** Distribution of mouth-hole contacts relative to the start of eating. **F.** Distribution of SH probabilities across individuals for each hole size (linear regression, r^2^ = 0.34, p-value < 0.001). **G.** Probability of Eat (teal) and SH (yellow) given that a mouth-hole contact was observed (p-value < 0.001, two-sided Fisher’s exact test). **H.** GLMM with subjects as random effect (mouth-hole contact: coefficient = –0.278, p-value < 0.001, other behaviors are not significant). **I.** 2D map of clamping locations of bottom (light blue) and top (light green) teeth for a peanut presenting a hole (black circle), red circle represents the clamp that is a teeth-hole contact, numbers correspond to the order of the clamp in the sequence. **J.** Distribution of movement vector orientations compared to a null distribution (top, *χ*2 test: p-value < 0.001, post-hoc bin analysis: [81°-90°]: p-value < 0.001, [45°-54°]: p-value < 0.01, others are not significant) and transition probabilities graph for longitudinal and transversal movements (bottom). **K.** Percentage of 3-movement motifs right before the first mouth-hole contact (L: longitudinal, T: transversal) and schematic showing what an LLT consists of. **L.** Average amplitude of T movements relative to first mouth-hole contact (average ± std). **M.** Schematic showing a potential mechanism for lip-mediated hole detection followed by a long transversal movement to breach the nut with the bottom teeth. **N.** Basic formulas describing the drift diffusion model (DDM). **O.** SH probability across hole sizes measured empirically and for the best fit DDM. **P.** Number of jaw clamps across hole sizes measured empirically and for the best fit DDM. **Q-R.** 20 simulated temporal trajectories of decision variables of the best fit DDM (**Q.** intact, **R.** 1 cm hole). *: p-value < 0.05, **: p-value < 0.01, ***: p-value < 0.001, no bracket: not significant.

Interestingly, when we quantified navigation on peanuts containing a 1 cm hole (N = 26), we noticed that the probability of T → T movements (self-transition of transversal movements) doubled between intact and breached peanuts – going from 0.26 (***Fig. 3I***) to 0.54 (***Fig. 4I, J bottom***). In contrast, L → L transition probability remained unchanged at 0.65 and 0.67 (**Fig. 3I, 4J bottom**). Remarkably, this T → T probability increase occurred immediately before mouth-hole contact (***Fig. S5D***), suggesting that agoutis transiently modify their search behavior only at the moment of hole detection. Consistent with this interpretation, 40% of hole detections were preceded by a stereotyped sequence consisting of one longitudinal movement followed by two transverse movements (an LTT motif; ***Fig. 4K***). Furthermore, the amplitudes of these final two transverse movements increased sharply immediately before contact, reaching approximately the geodesic distance required to move between the two faces of the peanut (corresponding to a ∼90° rotation; ***Fig. 4L***). In contrast, longitudinal movement amplitudes remained constant throughout shell assessment (***Fig. S5E***), indicating that only the transverse component of the search is selectively adjusted. Finally, we found that holes were contacted directly with the lower incisors in 83% of detection events (***Fig. S5F***). Together, these observations suggest that agoutis retain the same global search algorithm regardless of whether a hole is present, but refine their movements immediately before confirmation. We propose that agoutis first detect the edge of a hole — potentially using tactile information from the lips — and subsequently execute one additional long transversal movement to position the lower incisors over the opening (***Fig. 4M***), allowing direct tooth contact which serves as a confirmatory sensory event and triggers the decision to consume the peanut.

Behavioral scoring from observers provided highly compelling evidence for the role of mouth to hole contacts in underlying the decision to eat or scatter-hoard. To confirm this in a fully agnostic way we asked whether this decision implementation could be modeled by a drift diffusion model (DDM). This type of model is a widely used framework in neuroscience to describe evidence accumulation during decision-making under sequential sensory sampling(*45–49*). We modeled a decision variable (x_t,_ ***Fig. 4N***) that evolves over successive jaw clamps, drifting at a constant rate (*μ*; ***Fig. 4N***) toward a scatter-hoarding boundary (B_SH_, ***Fig. 4N***) representing the agoutis’ strong drive to SH during SH-mode. Each simulated jaw clamp was performed on a full-scale three-dimensional model of a peanut, allowing us to determine whether the lower teeth contacted a hole (contact, C = 1) or intact shell (C = 0; ***Fig. 4N***). A tooth-hole contact provided discrete evidence of shell damage by producing an instantaneous change in the decision variable (Δ; unconstrained in sign), potentially driving the decision variable across the eating boundary (B_Eat_, ***Fig. 4N***). To account for imperfect sensory processing, the model also allowed false-positive hole detections. We fit this realistic 3D model to our parametric data using the joint likelihood of our empirical SH probabilities (N = 11 individuals, ***Fig. 4O***) and number of clamps (***Fig. 4P, S5I***) across hole sizes; and we found that our DDM model can reproduce agouti decision making (***Fig. 4O-P***). We further analyzed the behavior of the decision variable x in time for different hole sizes (***Fig. 4Q-R, S5G-H***) as well as the value of the fitted parameters. Strikingly, the fitted jump magnitude was Δ = 0.52 ± 0.01, whereas the average separation between the scatter-hoarding and eating boundaries was only 0.83 units (B_SH_ = 0.31 ± 0.01, B_Eat_ = –0.53 ± 0.01). Thus, a single tooth-hole contact changed the decision variable by approximately 61% of the entire decision space. By comparison, the gradual drift during sampling was much smaller (μ = 0.10 ± 0.00), meaning that the evidence provided by one tooth-hole contact was equivalent to roughly five jaw clamps without contact.

Together, these results provide a computational explanation for our behavioral observations: shell assessment consists of gradual evidence accumulation during active exploration, while tooth-hole contact acts as a high-confidence sensory event that rapidly terminates the search and commits the animal to eating. This implementation naturally links the mechanics of shell exploration to a neurobiologically plausible evidence-accumulation process(*47*, *49*, *50*).

## Discussion

How foragers make decisions is of great importance for survival(*51*, *52*), reproduction(*53*, *54*), and ecosystem health(*1*, *55*, *56*). However, in most cases only the outcomes of foraging decisions are studied, with little insight into the computations, algorithms and implementations that drive them(*30–32*, *35*). In the present study we focused on the critical moment where agoutis make the decision to consume food items immediately or store them for later use. We discovered that agoutis engage in an active information-seeking assessment of preservability.

First, we revealed that agoutis transition into a persistent “scatter-hoarding mode” after multiple eating events, characterized by prolific caching and the emergence of oro-manual information-seeking behaviors. Once in this state, agoutis actively assess food items before deciding whether they are suitable for storage. Remarkably, agoutis readily scatter-hoarded empty peanuts with intact shells while consuming value-containing peanuts with compromised shells, showing that shell integrity, which promotes preservability, outweighed content value in determining whether a peanut should be cached or not. Second, we showed that preservability assessment is an efficient oro-manual information-seeking behavior. Agoutis navigate peanut shells by relying on cycles of rotations and clamping, producing sampling movements that exhibit an asymmetry that conforms to the anisotropy of the object they are manipulating and makes their search for holes optimal. This active sampling strategy enables agoutis to accumulate evidence of shell integrity into a decision threshold where hole detection, through mouth-hole contact, emerges as the strongest driver of consumption. Together, these findings suggest that agoutis use active sensing and integrate the 3D structure of the peanuts they manipulate to make long-term decisions estimating their preservability, an affordance that is key for food storage.

Previous studies have observed information seeking behaviors in wild squirrels(*57*). Preston & Jacobs have shown that fox squirrels perform head flicks that are correlated with nut species, complete absence of shell, and decision to cache. However, our study is, to our knowledge, the first account of a scatter-hoarding species that makes decisions based on haptic assessment of the external properties of food items(*29*, *30*, *37*). Our results contrast with previous studies proposing that perceived internal value of cached items, often inferred from weight, is a primary determinant of scatter-hoarding decisions(*30*, *37*).

A few studies have suggested rodents are capable of perceiving the perishability of different cached items(*56*, *58*, *59*). Interestingly, Jansen et al. have shown that agoutis deflesh *Astrocaryum* nuts before caching them, which reduces pilferage by making them less conspicuous(*25*). Our results strongly suggest that agoutis have evolved this cognitive capacity and emphasize the importance of preservability over immediate value, making it a great animal model for intertemporal choice(*17*), a fascinating and rare cognitive capacity(*60*) whereby an individual compares and actions and their resulting rewards when they occur at remote points in time.

Our work implies that future lab-based paradigms on information-seeking behaviors should consider the role of changing internal states(*22*, *61*) as well as integrate their ecologically relevant motivations. In the wild, future work will integrate models of value learning, optimal foraging, and temporal discounting to our understanding of scatter-hoarding intertemporal decision-making. While our findings highlight the mechanisms of scatter-hoarding decision-making with non-autochthonous nuts, it is important to acknowledge that other factors influence this process (variables such as nut type, hunger state, local food availability, competitor presence, and perceived risks of predation or pilferage have all been shown to affect caching behavior(*27*, *36*, *62–64*)). Most of these factors vary across our experiments, yet we observed highly consistent results that strongly follow shell integrity variations. This suggests the importance of preservability in scatter-hoarding decisions.

We have shown that agoutis are able to uncover breached regions of a peanut shell in just a few strategic rotations. These sophisticated manipulation capabilities rely on dexterous oro-manual coordination. This observation is in line with the evolution of thumb nails in rodents(*65*), which has been associated with diets requiring improved object handling. In fact, while eating peanuts, agoutis were observed systematically peeling even the internal skin of the peanuts with exquisite detail (***Fig. S5J***). These behavioral capabilities are likely supported by a large cortical expansion of their mouth somatosensory representation (lips, teeth, nose)(*38*, *66*), situated in close cortical proximity to their forelimb representation(*38*, *66–68*). Furthermore, we showed that agoutis make decisions based on the affordances conferred on fruits by their less nutritive yet protective shells. Like most frugivores they infer the value of objects with diverse combinations of endocarp, mesocarp and exocarp through a process termed extractive foraging(*69*, *70*) which has been linked to the intelligence of primates and birds(*69*, *71*, *72*). Altogether, these capacities make agoutis an extraordinary rodent model for the study of active sensing, sensorimotor intelligence, and affordances. Food manipulation is rare in neuroscience, however, studies of rodents manipulating artificial(*73*, *74*) or natural food sources(*75*, *76*) has the potential to uncover natural forms of information-seeking behavior(*61*, *77*).

Remarkably, the surface scanning and teeth-clamping of the agouti mirror human and primate gaze patterns during visual search(*78–80*). During visual foraging humans search, compare objects, and will fixate on the item they will select during value-based decision-making(*81*) analogous to the way agoutis explore peanut shells before finding the breaches and committing to eating or caching. Furthermore, in both cases asymmetries in the movements (clamping or gazing) adapt to anisotropies of the object or the scene being explored(*39*, *81*) (when a scene is long and horizontal, horizontal eye movements are favored). This suggests deep homologies between active search strategies which transcend species(*82*), dimensionality, or sensory modalities(*81*, *83*). The parallel between the agouti’s assessment capabilities and human gaze while searching a visual scene suggest that they may possess a rich representation of the 3D structure of objects, potentially supported by similar brain structures(*84*) and functional cell types(*85*).

## Supplementary figures

**Supplementary figure 1:**
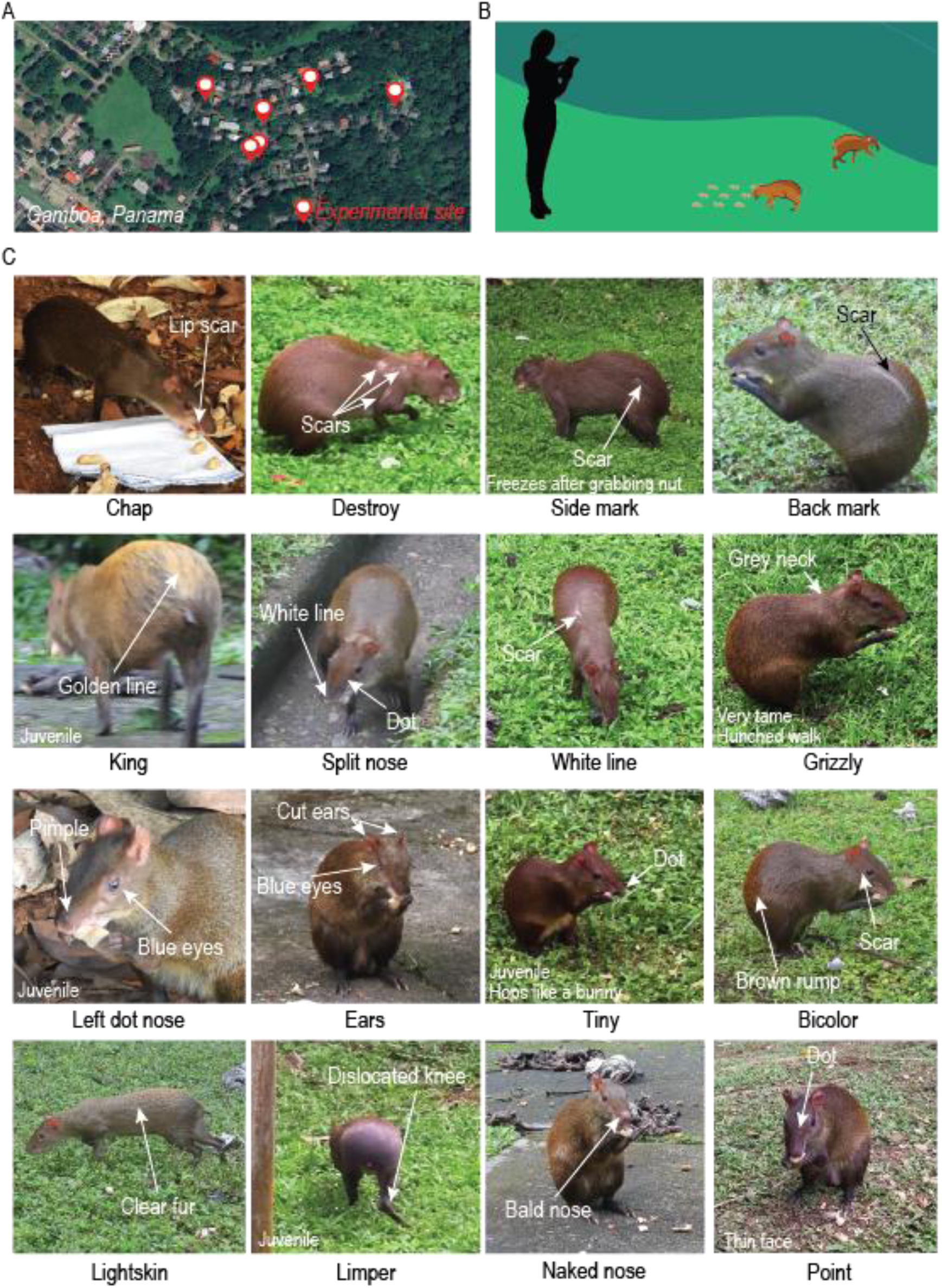
Field station and agouti visual markings. **A.** Experiments were run across 6 experimental sites in the field station in Gamboa, Panama. **B.** Wild agoutis freely engage in experiments by grabbing nuts of different conditions in their natural environment. **C.** 16 of the individuals that participated in experiments and examples of visual markings used to recognize them.

**Supplementary figure 2:**
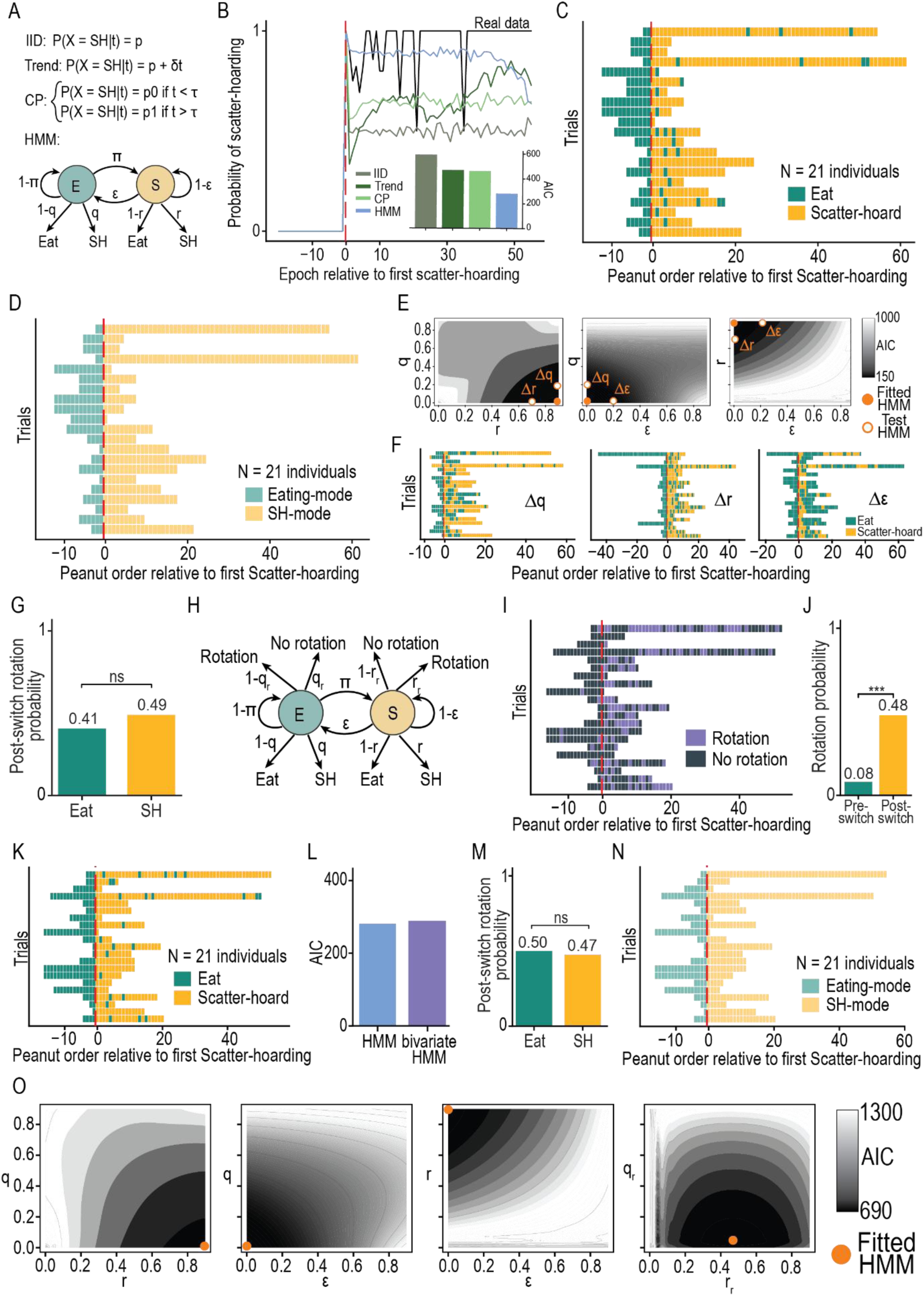
Agoutis switch to an absorbent SH-mode in which rotations emerge. **A.** Four models fitted to SH-mode experiment data: IID (Independent Identically distributed, random), Trend, CP (global Change Point), HMM (Hidden Markov Model). **B.** Probability of scatter-hoarding across epochs relative to first SH instance for empirical data and four models. Inset: AIC score for the fit of the four models (AIC: HMM = 281, CP = 461, Trend = 468, IID = 591). **C.** Emissions of the best fitted HMM model. **D.** State distribution for the best fitted HMM model. **E.** AIC score landscapes for the best fitted HMM and three test HMMs (Δq: fitted q + 0.2, Δr: fitted r + 0.2, Δε: fitted ε + 0.2) in spaces (q,r), (q, ε), and (r, ε). **F.** Emissions of behaviors for the three test HMMs (Δq, Δr, Δε). **G.** Probability of Eat and SH after switching to SH-mode in empirical data (p-value = 0.46, two-proportion Z-test). **H.** Bivariate HMM emitting behaviors and rotations. **I.** Rotation emissions for the best fitted bivariate HMM. **J.** Rotation probability pre– and post-switch to SH-mode for the best fitted bivariate HMM (p-value < 0.001, independent two-sample Welch’s t-test). **K.** Behavior emissions for the best fitted bivariate HMM. **L.** AIC score for the best fitted HMM and bivariate HMM fitted to empirical behaviors. **M.** Probability of Eat and SH after switching to SH-mode in empirical data (p-value = 0.81, independent two-sample Welch’s t-test). **N.** State distribution for the best fitted bivariate HMM model. **O.** AIC score landscapes for the best fitted bivariate HMM in spaces (q,r), (q, ε), (r, ε), and (q_r_, r_r_).

**Supplementary figure 3:**
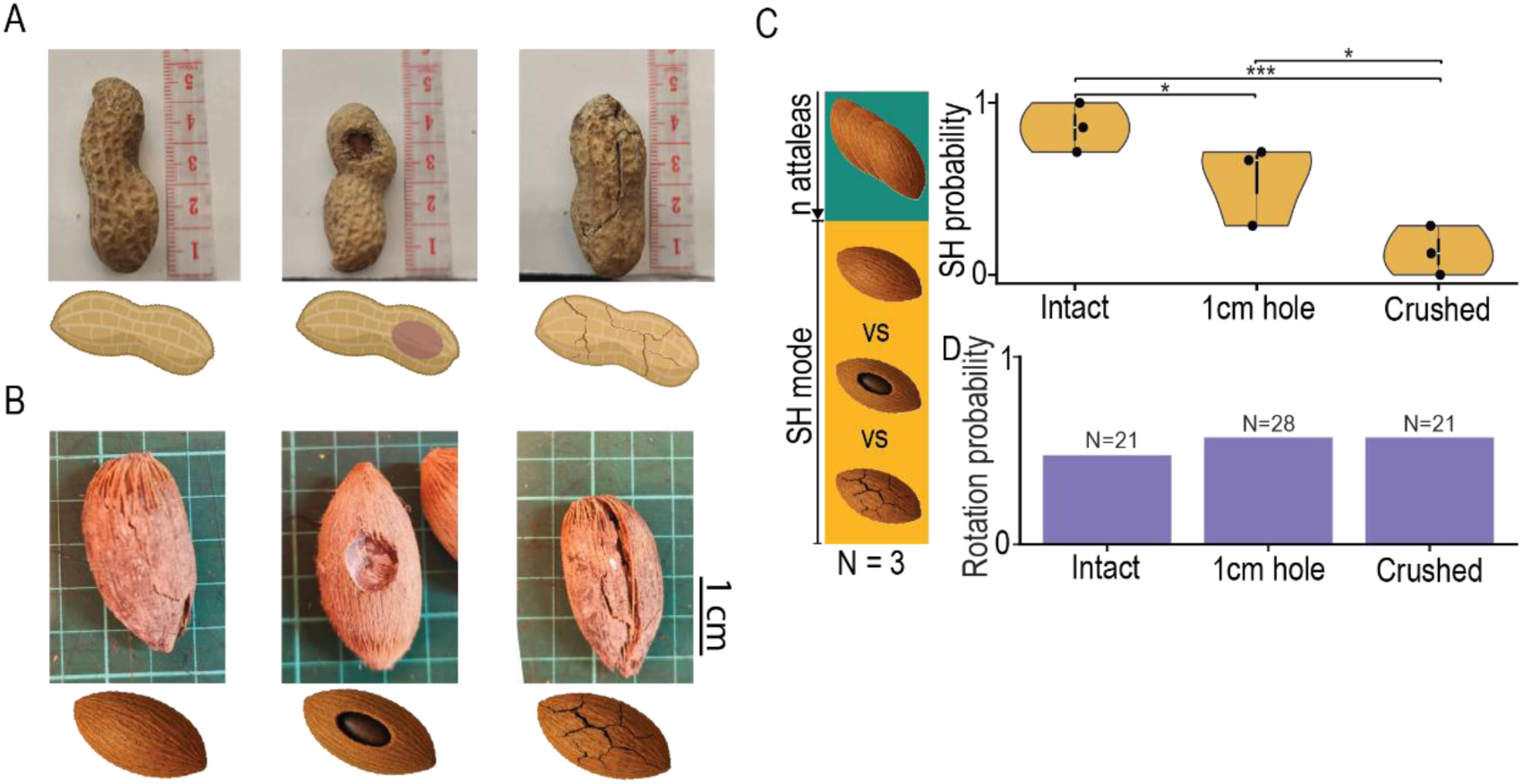
Experimental nuts and scatter hoarding results with *Attalea butyracea*. **A**. Peanuts of three conditions: intact (left), 1cm hole (middle), crushed (right). **B.** *Attalea butyracea* nuts of three conditions: intact (left), 1cm hole (middle), crushed (right). **C.** Experimental procedure to test integrity decision making in naturally-occurring *Attalea* seeds.

**Supplementary figure 4:**
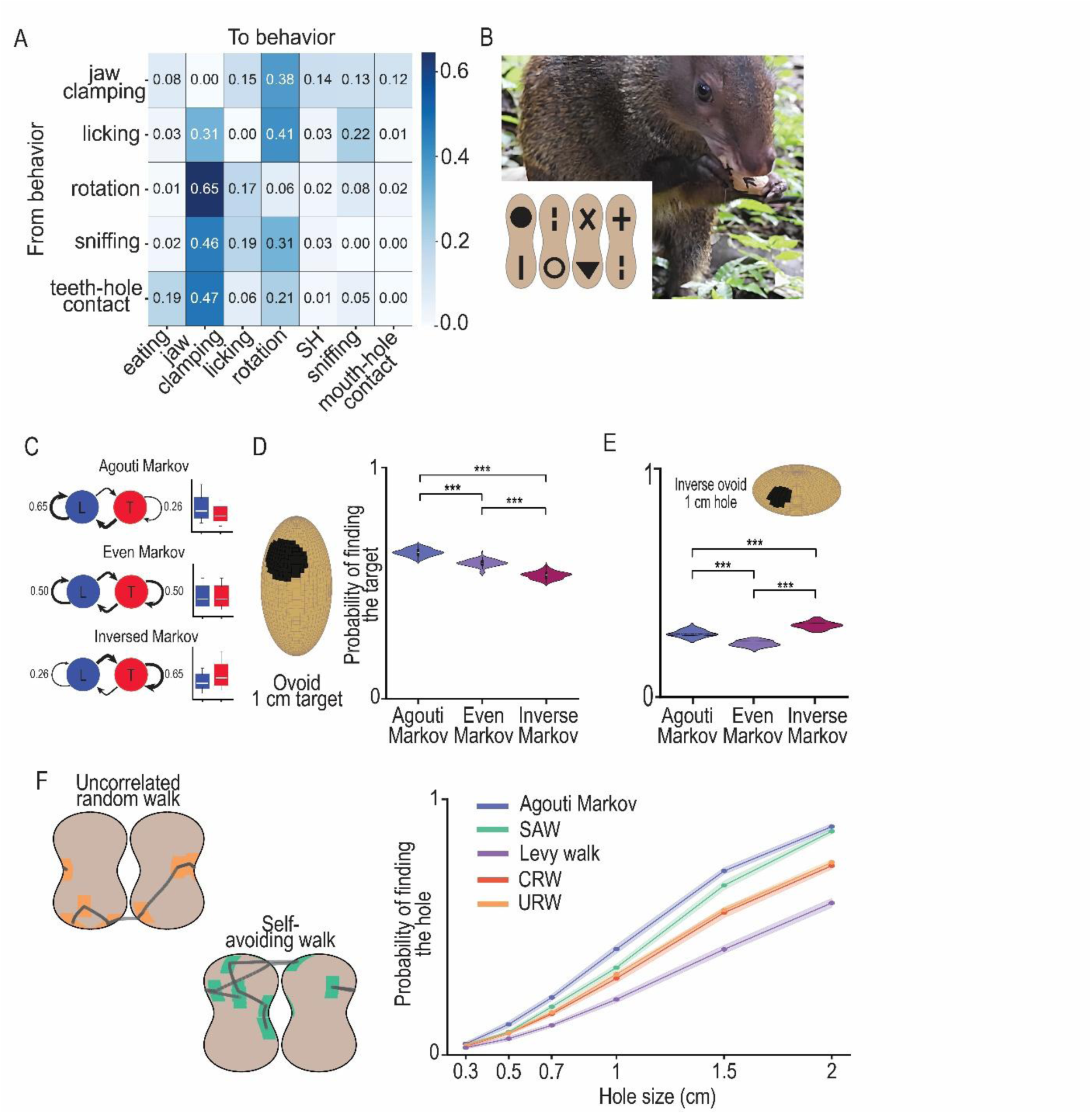
Ethogram for parametric experiment, peanut visual landmarks and inverse ovoid simulation. **A.** Transition probability matrix of assessment behaviors. **B.** Example of agouti manipulating a peanut with artificial visual landmarks. **C.** Transition probabilities (right) and vector length distributions (left) for data-driven Markov walk (top), Even Markov (middle, no asymmetry), Inversed Markov (bottom, inverse asymmetry). **D-E.** Performance of Agouti Markov, Even Markov, and Inversed Markov walks at finding a 1cm diameter target on **D.** 3D ovoid shape (peanut anisotropy) **E.** 3D inverse ovoid shape. **F.** Uncorrelated Random walk (URW, orange) and Self-Avoiding Walk with memory 1 (SAW, green) sampling the surface of the peanut shell using clamps (colored rectangles) and probability of clamp-hole contact across 10,000 simulations for each type of walk.

**Supplementary figure 5:**
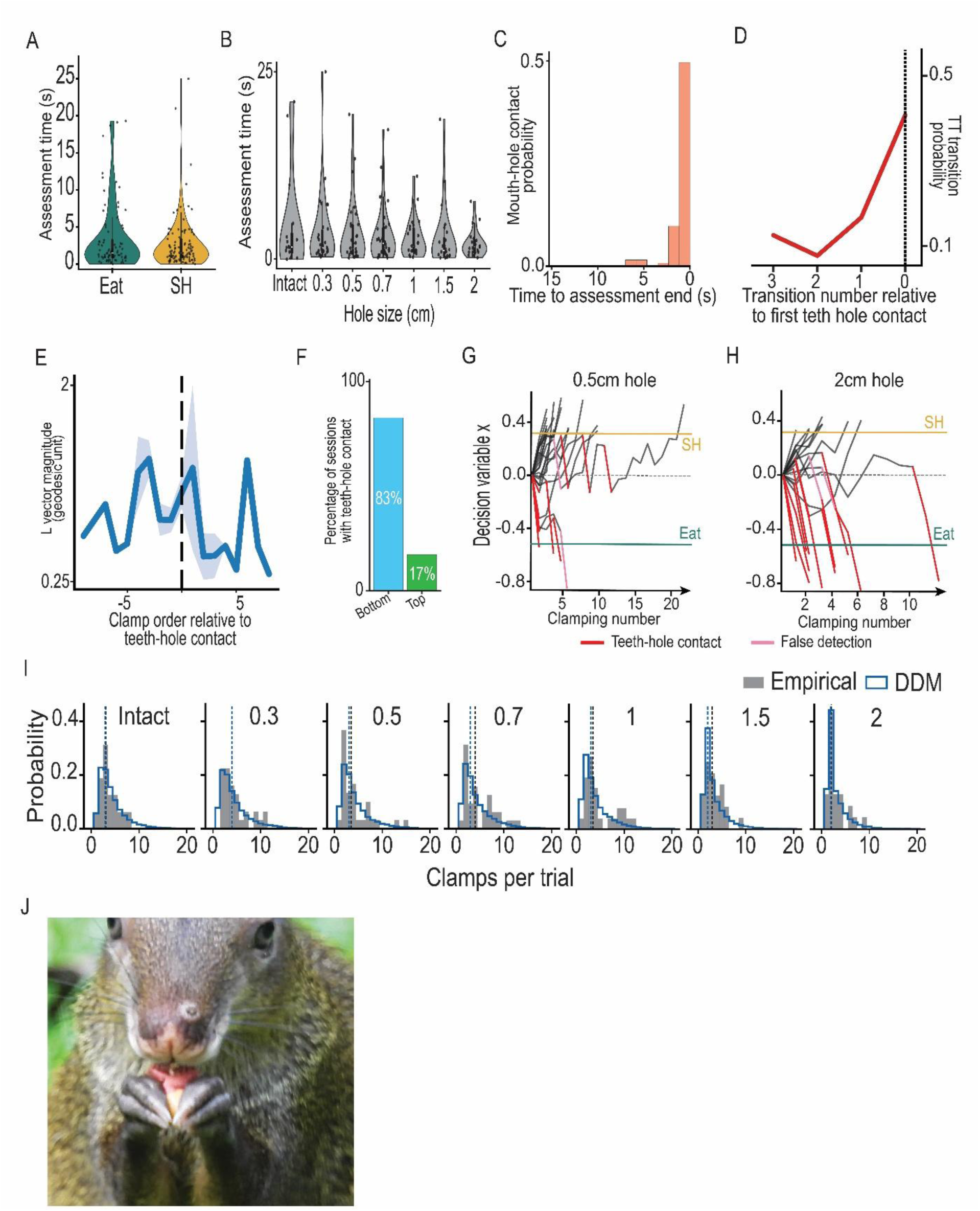
Further descriptions of assessment statistics. **A.** Distribution of assessment time (seconds) for peanuts that were eaten or scatter-hoarded. **B.** Distribution of assessment time per hole sizes. **C.** Mouth-hole contact probability throughout assessment duration. **D.** T → T transition probabilities relative to first teeth-hole contact (transition 0). **E.** Average magnitude of longitudinal vectors across clamps relative to first teeth hole contact. **F.** Share of trials where bottom (light blue) or top (light green) teeth perform mouth contact. **G-H.** 20 simulated temporal trajectories of decision variables of the best fit DDM (**G.** 0.5 cm hole, **H.** 2 cm hole). **I.** Comparison of probability distributions of clamp numbers per trials between empirically measured data and best fitted DDM for intact and breached peanuts of all hole sizes. **J.** Example of an agouti peeling the skin of a peanut seed.

## Supplementary Movies

**Supplementary video 1: Agouti scatter-hoarding**.

Link: https://doi.org/10.6084/m9.figshare.33307836

**Supplementary video 2: Extensive peanut rotation during assessment of the agouti**.

Link: https://doi.org/10.6084/m9.figshare.33307854

**Supplementary video 3: Agouti assessing a peanut before making a scatter-hoarding decision**.

Link: https://doi.org/10.6084/m9.figshare.33307872

**Supplementary video 4: 3D tracking of peanut rotation for integrity assessment by an agouti**.

Link: https://doi.org/10.6084/m9.figshare.33307881

**Supplementary video 5: Teeth-hole contact during integrity assessment in the agouti**

Link: https://doi.org/10.6084/m9.figshare.33307899

## Acknowledgements

We thank Andy Quitmeyer and Kit Claro for their precious help in understanding and learning about agoutis. We thank Yossi Yovel, Saikat Ray, Lyle Kingsbury, Adam Lowet, Kristian Herrera, Andrew Bahle, Kelsey Tyssowski and Michael Brecht for comments on the initial and subsequent drafts of the manuscript. We thank Rachel Page, Bill Wieslo, Owen McMillan, and Isis Ochoa from the Smithsonian Tropical Research Institute for their ongoing support as well as the Sistema Nacional de Investigacion, Association de Interes Publico (SNI AIP). This manuscript is dedicated to the memory of the ever generous and inspiring Adam Kampff.

## Author Contributions

Conceptualization: LD, JISS

Methodology: LD, ET, MW, DG, CT, JISS

Experimental work: LD, ET, MW, CT, DG, JISS

Behavioral Scoring: MW, ET, LD

Data Analysis: LD, ET, JISS

Coding: LD

Visualization: LD, MW, JISS

Funding acquisition: JISS

Project administration: JISS

Supervision: JISS

Writing-original draft: LD, CT, DG, JISS

Writing-review and editing: LD, ET, MW, DG, CT, JISS

## Data, code, and materials availability

https://github.com/Lucas97223/Oro-manual-assessment-of-food-storage-stability-in-a-wild-rodent

## Materials and Methods

### Ethical Note

This study was conducted under the permit of Ministerio del Ambiente, Panamá (SE/A-96-15, SE/A-96-18, SE/A-38-2020), and the Animal Care and Use Committee at the Smithsonian Tropical Research Institute (SI-24058, SI-23061) and adhered to the ASAB/ABS Guidelines for the Use of Animals in Research. No invasive interventions were used, and participation was entirely voluntary.

### Study site and study population

The study was conducted in the wild at the Smithsonian Tropical Research Institute (STRI) scientific station in Gamboa, Panama – over two field seasons (6 weeks – March 21st to May 21st 2025 – and 13 weeks – December 8th to March 8th 2026). The target species was the Central American agouti (*Dasyprocta punctata*), a diurnal scatter-hoarding rodent. Observations were made under natural foraging conditions within the animals’ territories. Food items were placed in natural open areas used frequently by agoutis. Wild agoutis in this area are habituated to the presence of humans and can be observed at close distances. Any agouti participating in an experiment is fully wild and was engaging in the task on its own and could leave the experiment whenever it wanted. Each agouti participating in an experiment was visually identified using morphology or specific physical traits.

### Study design and Procedures

#### 1. Fruit type and behavioral action sequence

*Protocol —* Here we aimed to characterize the fruit-dependent foraging ethogram in wild agoutis, focusing on the behavioral sequences and decision-making strategies involved in fruit evaluation, dissection, consumption, and hoarding. To document naturalistic, value-based foraging decisions, we constructed baited feeding arenas in the field, each measuring 3 × 3 fruit positions (9 fruits per trial). Each trial presented three whole fruits from each of three distinct types, selected from a pool including banana, grape, apple, orange, and peanut. The spatial arrangement of fruits was pseudo-randomized across trials to prevent location bias. Since individual identification was not performed for this experiment, we considered that each fruit was treated independently.

*Data collection and behavioral measurements* — Two experimental sessions were conducted daily—one between 07:00 and 09:00 and the other between 16:00 and 18:00—corresponding to peak periods of agouti activity. During each session, multiple foraging arenas were simultaneously deployed across different locations. Each feeding arena was monitored using a multi-angle video setup consisting of three fixed cameras positioned around the arena to maximize visibility and ensure full coverage of fruit interaction sequences. Trials were conducted under varying environmental conditions, with systematic logging of metadata including date, time of day, temperature, weather, and study site. Each animal-fruit interaction was characterized by a sequence of different behaviors, categorized into non mutually exclusive categories as described in Table 1.

**Table 1.** Ethogram of agouti’s foraging behavior.

| Behavior | Description | Scoring |
| --- | --- | --- |
| Sniffing | Bringing its nose into close proximity with the fruit to inspect it olfactorily, without physical contact beyond sniffing. | Binary (0/1) |
| Rotation | Turning or manipulating the fruit along its axis by the forelimbs or mouth, typically while being held. | Binary (0/1) |
| Transporting | Engaging in active locomotion with the fruit in the mouth, relocating it to a new spot for immediate eating. | Binary (0/1) |
| Eating | Masticating or consuming food items. | Binary (0/1) |
| Scatter-hoarding | Sequential behavior in which the animal carries a food item in its mouth to a chosen site with suitable substrate, digs a small pit using its forelimbs, drops the item into the hole, presses it down firmly, refills the pit with soil using forward strokes of its forelegs, and often covers the spot with leaves or other debris. | Binary (0/1) |

#### 2. Scatter-hoarding (SH) experiment

*Nuts used —* two types of nuts were used throughout all the experiments presented in this study. Most experiments were run using peanuts found in local stores. Peanuts are not a native nut in the agouti’s natural environment. However, they are very attractive to agoutis and allows us to have wild individuals that engage in our experiments. Furthermore, peanuts are rather small – their size is equivalent to most nuts agoutis find in their environment – which gives us the possibility to observe natural nut manipulation from agoutis. Finally, peanut shells are easy to break into, which makes experiments faster and removes any experimental confound that would be related to the effort to access the nut’s inside. Different peanut conditions are mentioned throughout our experiments:

– control peanut: two-chamber, symmetric peanut with a fully intact shell (no hole or breach, no irregularities).
– empty peanut: control peanut that was surgically opened on the suture line of the shell. The cotyledons of the peanuts are removed and the two half-shells are glued back together using a thin filament of hot glue. We acknowledge that the filament of hot glue might disturb the regularity of the peanut shell, though its thinness is mostly imperceptible from the outside.
– peanut with holes: control peanut in the shell of which a hole was carved using a pointy knife and a tiny pair of scissors. The hole position, on the biggest chamber of the peanut, is constant regardless of the size of the hole, measured using a meter.
– crushed peanut: control peanut which shell was manually compressed until it presented cracks and breaches uniformly on all its surface. Crushed peanuts present many thin cracks all over their shells that are not opened into holes, they conserve their full composure: they can be handheld without breaking down into pieces

The other type of nut that was used is nuts from the fruit of the palm tree *Attalea butyracea* that are harvested by hand on the rainforest floor, in the agouti’s natural environment.

*Protocol —* in all experiments the subjects were wild agoutis that were roaming in different locations consistently chosen at the edge of the rainforest at our field station in Gamboa. The individuals are free to engage with the experimenter and perform the experiment and are free to leave the experiment at any time. Experiments are run in the ecological niche of the individuals where no disruption is done to the environment, and within their natural territory. Predators were never observed at the times and locations where experiments are performed. Generally numerous conspecifics are present during the experiment, potentially disturbing the subject’s behavior, they are led further away from the experiment when possible. Overall, these experiments reflect the decision-making mechanisms that agoutis make in their natural environment, embedded in its complexity and variability at different timescales. All experiments mentioned below were run after the focal agouti of the experiment was brought to SH-mode. Each observation session began with the *ad libitum* provisioning of intact peanuts, unless mention of the contrary. Once an individual performed three consecutive scatter-hoarding acts of intact peanuts, that animal was designated as a focal agouti for the remainder of the session. Due to observational constraints, 1 to 4 agoutis were selected per session as focal individuals, depending on visibility and tracking feasibility.

##### a. Ethogram

Sessions were video-recorded using GoPro cameras or a professional video camera, following the focal agouti.

##### b. Behavioral scoring

Scoring of all the behaviors defined in the ethogram (***Table 2***) was done by one human observer on video recordings of the animals. Videos where it is impossible to detect what the animal is doing for all or part of the video are discarded from the analysis. The only exception is for the scoring of the outcome of the decision: agoutis eat in place or move a little bit if they are bothered by a conspecific but always stay in the frame of the camera; however one can clearly see that an agouti has decided to scatter-hoard when they leave right away after assessment and disappear in the bushes. In the latter context, the video is kept even though the animal disappears and it is scored as ‘scatter hoarding’. Mouth/Teeth-hole contacts are trickier to detect in videos, hence two different observers annotated this specific behavior. One of the observers was agnostic to the outcome of the assessment (i.e. decision to eat or scatter hoard).

**Table 2.**
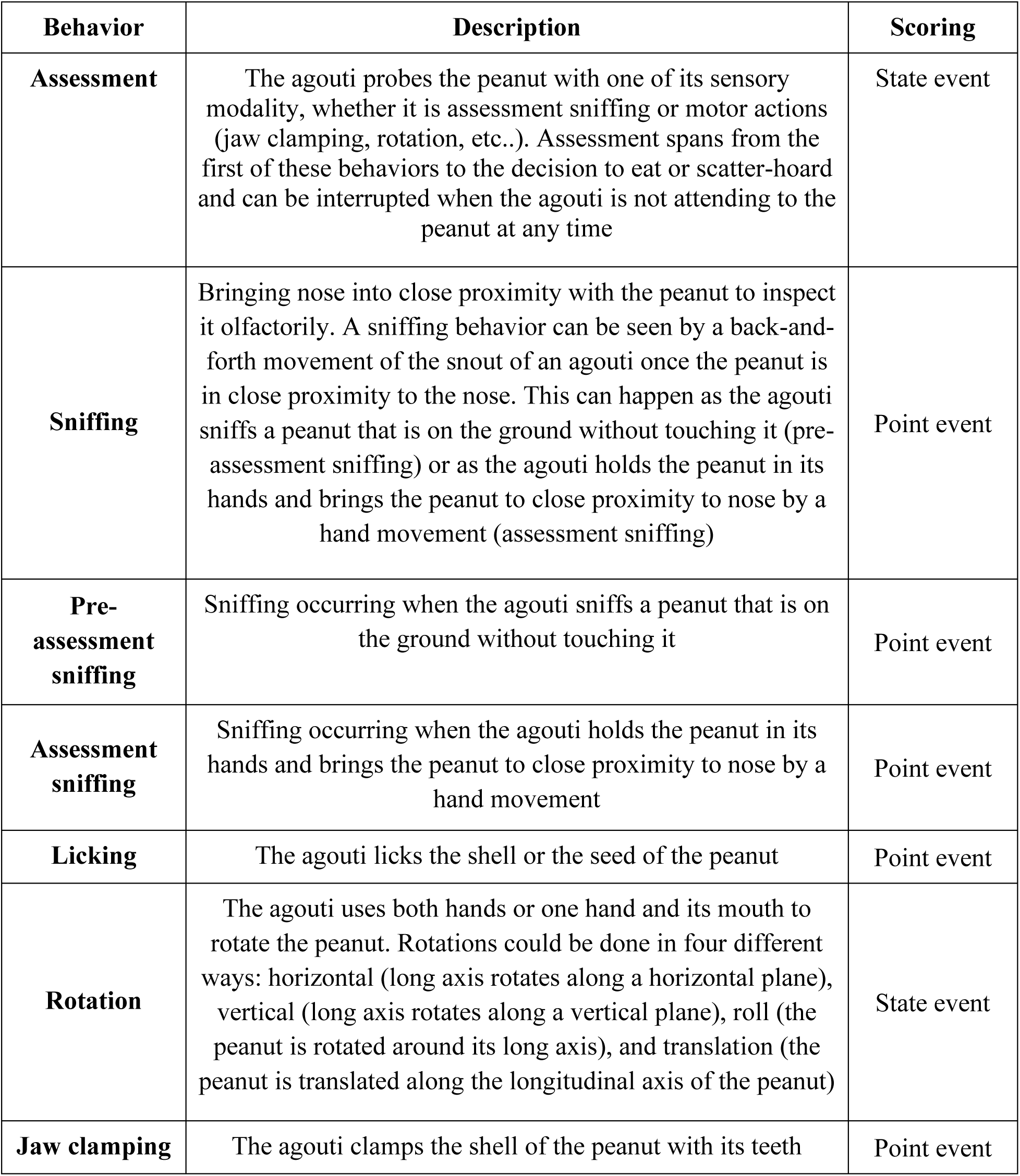
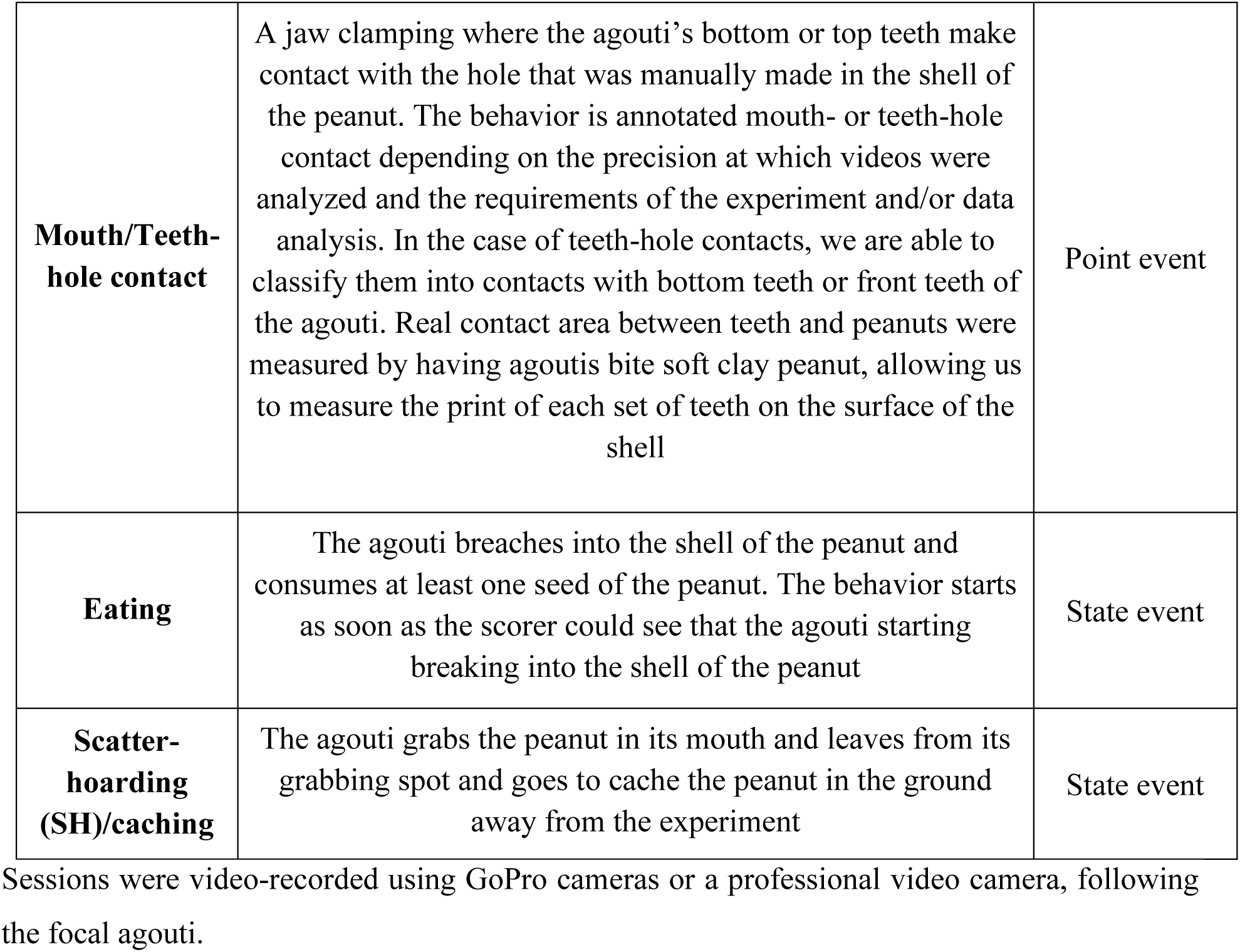
Ethogram of agouti’s peanut assessment and decision-making.

| **Behavior** | **Description** | **Scoring** |
| --- | --- | --- |
| **Assessment** | The agouti probes the peanut with one of its sensory modality, whether it is assessment sniffing or motor actions (jaw clamping, rotation, etc..). Assessment spans from the first of these behaviors to the decision to eat or scatter-hoard and can be interrupted when the agouti is not attending to the peanut at any time | State event |
| **Sniffing** | Bringing nose into close proximity with the peanut to inspect it olfactorily. A sniffing behavior can be seen by a back-and-forth movement of the snout of an agouti once the peanut is in close proximity to the nose. This can happen as the agouti sniffs a peanut that is on the ground without touching it (pre-assessment sniffing) or as the agouti holds the peanut in its hands and brings the peanut to close proximity to nose by a hand movement (assessment sniffing) | Point event |
| **Pre-assessment sniffing** | Sniffing occurring when the agouti sniffs a peanut that is on the ground without touching it | Point event |
| **Assessment sniffing** | Sniffing occurring when the agouti holds the peanut in its hands and brings the peanut to close proximity to nose by a hand movement | Point event |
| **Licking** | The agouti licks the shell or the seed of the peanut | Point event |
| **Rotation** | The agouti uses both hands or one hand and its mouth to rotate the peanut. Rotations could be done in four different ways: horizontal (long axis rotates along a horizontal plane), vertical (long axis rotates along a vertical plane), roll (the peanut is rotated around its long axis), and translation (the peanut is translated along the longitudinal axis of the peanut) | State event |
| **Jaw clamping** | The agouti clamps the shell of the peanut with its teeth | Point event |
| **Mouth/Teeth-hole contact** | A jaw clamping where the agouti's bottom or top teeth make contact with the hole that was manually made in the shell of the peanut. The behavior is annotated mouth- or teeth-hole contact depending on the precision at which videos were analyzed and the requirements of the experiment and/or data analysis. In the case of teeth-hole contacts, we are able to classify them into contacts with bottom teeth or front teeth of the agouti. Real contact area between teeth and peanuts were measured by having agoutis bite soft clay peanut, allowing us to measure the print of each set of teeth on the surface of the shell | Point event |
| **Eating** | The agouti breaches into the shell of the peanut and consumes at least one seed of the peanut. The behavior starts as soon as the scorer could see that the agouti starting breaking into the shell of the peanut | State event |
| **Scatter-hoarding (SH)/caching** | The agouti grabs the peanut in its mouth and leaves from its grabbing spot and goes to cache the peanut in the ground away from the experiment | State event |

Sessions were video-recorded using GoPro cameras or a professional video camera, following the focal agouti.

##### c. Scatter-hoarding mode experiment

In order to investigate the temporal structure of scatter-hoarding behavior, we gave a series of intact peanuts to single focal agoutis by throwing the nuts in front of them as they approached the experimenter. Observers scored the outcome of the decision, coding whether the peanut was eaten or cached by the agouti as well as whether a rotation of the peanut was performed before committing to their decision.

##### d. Internal nutritional content experiment

In order to test whether rotation behavior is used to evaluate the internal content of the peanut – such as weight or sound – single focal agoutis were presented with peanuts under two different internal content conditions: intact peanuts (intact, edible contents) and empty peanuts (shells reassembled, no contents). Each individual was presented with a pseudo-random mix of the two conditions during naturalistic foraging on open ground, where they would throw peanuts one at a time. 50 peanuts of each condition were weighed individual on a precision scale to study their weight distribution.

##### e. External shell integrity experiment

In order to test whether rotation was used to evaluate external shell integrity – such as damage or structural weakness – single focal agoutis were presented with peanuts under multiple different external conditions, organized in an arena format. An arena was fully deployed on the ground, prior to focal agouti approach. Each arena consisted of a 4×2 grid (8 peanuts per trial) with four experimental peanut shell conditions: intact (unmodified peanut shells); shells with a 1cm hole; shells with a 2cm hole peanut, and crushed shells (cracked shells simulating structural weakness). The conditions were distributed equidistantly and pseudo-randomly across the grid to avoid positional bias. Each observation session was divided into blocks, where a block was defined as a sequence of one or more consecutive arenas presented to the same focal agouti, as long as it remained in view and engaged. This structure allowed multiple arenas to be presented per individual per session (preserving trial independence at the level of the arena). The same experimental procedure was run with the *Attalea butyracea* seeds that are native from agoutis’ ecological niche, with conditions intact, shells with 1cm hole and crushed shells, in a 3×3 grid per arena. Videos were recorded of every single peanut.

##### f. Parametrics experiment

In order to investigate the integrity assessment capabilities of agoutis, we performed a parametric experiment with a wider range of hole sizes (intact; 0.3cm; 0.5cm; 0.7cm; 1cm; 1.5cm; 2cm; 2.5cm) in a 3×3 arena setting (2 intact peanuts and one of each type). This allowed us to examine at a finer resolution the relationship between shell damage and scatter-hoarding behavior. Videos were recorded of every single peanut.

##### g. Peanut tracking

In order to perform precise tracking of the rotations of the peanuts as well as the locations of the jaw clamping from the agouti on the peanut shell we provided agoutis with intact peanuts whose shells were marked with 8 unique symbols that would allow us to orient the peanuts in space at any time. Videos were recorded of every single peanut. We created an app that supports a 3D model of a peanut at scale, oriented with these unique symbols, which allowed us to locate each clamping site with precise coordinates and therefore extract clamp-to-clamp vectors for further analyses. Vectors are classified into longitudinal and transversal ones based on the component (Z or Arc axis of the peanut) that as the highest magnitude. All distances and magnitudes of vectors that are placed on the 3D model of a peanut are given in geodesic distance in centimeters based on the dimensions of the peanut model itself (see below).

#### 3. Longevity of different conditions of peanuts in the ground

To assess whether the condition of peanut shells would impact their longevity, or long-term value, after being buried in the ground by agoutis, we manually buried peanuts in the ground during two experiments to get a sense of their differential perishabilities, as well as their differential survival to pilfering. In the first case, experimenters manually buried 357 peanuts (10 peanuts for turgidity and 7 peanuts for dry mass loss and 7 peanuts for molding analysis retrieved each day for 7 days). These peanuts were buried in the ground 4cm deep, which is the average caching depth of agoutis, and were protected by an iron grid so they could not be pilfered. Dry mass loss was measured with the following procedure: peanuts are retrieved from the ground, cleaned, seeds are extracted and weighed, they are then dried and weighed again; the difference between the mass after and the mass before is the dry mass loss. Molding analysis was done by first calculating the baseline color of pixels of healthy peanuts seed and then computing the average deviation from this color for each seed. With constant light settings, these differences in colors are explained by either black or white mold that develops on the seed of the peanut. Turgidity was measured as a proxy to humidity content. We would press on half peanut seeds with a pressure gauge and extract the force value (in N) that is necessary to break or tear the seed in at least two parts. In the second case, experimenters buried pairs of intact and crushed shell peanuts 4 cm deep and 20 cm apart in front of camera traps in different locations of our field station (gardens, forest, forest edge, trails). We scored which of the two peanuts was retrieved first as well as what species of animals did the retrieval (95% were agoutis, 5% were coatis).

#### 4. Computational modeling

##### a. SH-mode

To determine whether the transition from eating to scatter-hoarding represents a gradual shift in motivation or an abrupt behavioral state change, we modeled the sequential foraging decisions of the subjects (N = 21). For each animal, the foraging sequence was encoded as a discrete binary time series y(t) ∊ {0, 1} over trials t = 1, …, T, where y(t) = 0 denotes eating and y(t) = 1 denotes scatter-hoarding. We evaluated four nested probabilistic models against these sequences:

– Independent and Identically Distributed (IID) Model: Serving as a baseline, this model assumes the probability of scatter-hoarding remains constant throughout the foraging session, such that P(y(t) = 1) = p.
– Trend Model: To test for a gradual motivational shift, we modeled the probability of scatter-hoarding as a function of the normalized trial order, P(y(t) = 1) = p + δt, where δ dictates the steepness of the temporal trend.
– Global Change-Point (CP) Model: To test for an abrupt, step-like shift in behavior, we fitted a piecewise-constant probability model characterized by a single structural change-point at trial *τ*. The probability of scatter-hoarding was defined as p_0_ for t < *τ* and p_1_ t > *τ*.
– Hidden Markov Model (HMM): To capture underlying, unobservable behavioral states that dictate noisy emissions, we implemented a 2-state discrete HMM comprising an ‘Eating’ state (E) and a ‘Scatter-hoarding’ state (S). The transition dynamics were parameterized by *π* (probability of transitioning from E to S) and ε (probability of transitioning from S to E). The emission probabilities for observing a scatter-hoard given the true underlying state were defined as P(y(t) = 1 | E) = q and P(y(t) = 1 | S) = r.

To estimate the optimal parameters for all models, we maximized the joint log-likelihood pooled across all animal sequences. For the IID and Logistic Trend models, we utilized iteratively reweighted least squares (IRLS). For the CP model, we performed an exhaustive grid search to find the optimal global *τ*, p_0_, and p_1_ that maximized the pooled likelihood. For the HMM, parameters were estimated using the Expectation-Maximization (EM) algorithm (Baum-Welch algorithm). During the E-step, expected state counts (from the forward-backward algorithm) were pooled across all sequences to iteratively refine the global *π*, ε, q, and r values until convergence (tolerance of 10^-9^). To compare relative model performance while penalizing for model complexity, we computed the Akaike Information Criterion (AIC) for the global fit of each model.

To visually assess model fit, we performed predictive simulations drawing synthetic behavioral sequences from the fitted global parameters of each model. Both empirical and simulated sequences were aligned to the epoch of the first observed scatter-hoarding event, and the mean sequence probabilities and 95% predictive bands were extracted. For the HMM, to confirm the robustness of the fitted global minimum and visualize parameter identifiability, we conducted a grid-search to calculate the AIC landscape across two-dimensional parameter planes. Finally, to mechanistically validate the role of specific transition and emission parameters on sequence formation, we simulated alternative behavior sequences by systematically perturbing individual HMM parameters (Δq, Δr, Δε) relative to the optimal fit.

We also built a bivariate HMM that shared the same structure as the original HMM that was the best fit but added another type of emissions: rotation or no rotation. The emission probabilities of outcomes (eat or SH) were fitted to the observed behavior as explained before, the rotation emissions were fitted to the rotation observations in the same way. The best bivariate HMM was the one that maximized the joint likelihood of outcome and rotation emissions. Rotation and outcome emissions are fully independent. AIC landscapes are built in the exact same way as described above, adding an additional sweep of the two parameters that control rotation emission in Eating state (q_r_) and in SH state (r_r_).

##### b. 3D peanut model and ovoids

We mathematically defined the geometry of the seed substrate using a standardized 3D parameterized surface. Natural peanuts closely resemble a three-dimensional solid of revolution generated by a modified Cassini oval. This geometry provides a continuous, differentiable surface characterized by two distinct symmetric lobes separated by a central medial constriction (the “waist”), allowing for accurate spatial mapping and vector trajectory simulations across varying surface curvatures.

The seed substrate was modeled as a surface of revolution defined in a cylindrical coordinate system (r,θ,z), where z represents the primary longitudinal axis of the seed, r is the transverse radial distance from the longitudinal axis, and θ is the azimuthal angle. The outer surface of the seed is defined by the radial function r(z), which governs the seed’s profile: r(z) = max(0, b^4^+4a^2^z^2^−z^2^−a^2^). This equation is controlled by two strictly positive scale parameters, a and b, which dictate the geometric proportions of the solid. The parameter a controls the separation distance of the geometric foci (dictating the distance between the two lobes), while the parameter b governs the overall volume and thickness of the resulting lobes and the depth of the central constriction. To map this generalized equation to the specific morphological constraints of the natural seeds consumed by the subjects, empirical measurements of the seeds were extracted and optimized to yield the scale parameters. For all geometric and behavioral simulations, the model parameters were fixed to: a = 1.2602 cm and b = 1.3749 cm. This mathematical formulation naturally produces a peanut-like topology characterized by two outward bulging lobes separated by a central medial constriction (the “waist”). Based on the parameters, the model features a total longitudinal length of 3.73 cm, a maximal lobe diameter of 1.50 cm, and a pinched waist diameter of 1.10 cm. The continuous surface of the seed was discretized into a highly dense polygonal mesh for Markov chain Monte Carlo simulations. Behavioral tracking data—namely the discrete, orthogonal movements comprised of longitudinal advancements (along the z-axis) and transverse rotations (arcs along the azimuthal angle θ)—were projected directly onto this surface manifold. The accurate representation of surface curvature (from the 1.10 cm waist to the 1.50 cm lobes) ensures that rotational step magnitudes (arc lengths) appropriately deform based on the local radius r(z) of the seed at any given time step t. This dynamically and accurately reflects the animal’s physical traversal across a natural seed, accounting for the changing local diameter and allowing for precise spatiotemporal calculation of hole-finding probabilities.

The standard convex “Ovoid” (ellipsoid) and the “Inverted Ovoid” were both mathematically derived to ensure they share the exact same maximal boundaries and surface areas, allowing for direct comparative analysis without confounding overall size differences. To serve as a strictly convex baseline—eliminating the structural complexity of the waist—a standard Ovoid (ellipsoid of revolution) was constructed to perfectly encapsulate the Inverted Ovoid. To achieve this, the maximal dimensions of the Inverted Ovoid were extracted and mapped directly to the axes of the standard Ovoid: the semi-major (longitudinal) axis was set to match the absolute length of the natural seed (1.865 cm from origin to pole) and the semi-minor (transverse) axis was set to match the maximum radius of the seed’s lateral lobes (0.75 cm).

##### c. Sampling walks

To systematically evaluate the functional efficiency of the agouti’s search heuristic against canonical theoretical search models, spatial traversal simulations were conducted across a gradient of target (hole) sizes. The models were tested on the aforementioned continuous 3D Cassini oval manifold representing the natural geometry of the seed. The simulation framework quantified the terminal probability of successfully locating a centralized geometric target (a hole situated at the medial constriction, z=0, θ=0) within a strictly constrained energy budget (N=4 longitudinal/transverse movement steps, equivalent to 8 total jaw clamps).

Five distinct generative search strategies were subjected to the spatial simulation. Each strategy initiated from a random uniformly distributed coordinate on the seed’s surface and evolved through N steps according to distinct navigational rules:

– Empirical Markov (Agouti) Model: This model replicates the observed, highly polarized kinematic behavior of the agouti. The movement state (longitudinal vs. transverse) evolved via the empirically derived transition matrix (p(LL) =0.65, p(TT) = 0.26). Step magnitudes were stochastically drawn from bounded normal distributions fitted to empirical kinematics (Longitudinal μ=0.967 cm; Transverse μ=0.864 cm, with σ=0.3). Longitudinal directionality was biased toward the seed’s poles according to the animal’s empirical spatial preferences, while transverse directionality remained strictly unbiased.
– Self-Avoiding Walk (SAW, M=1): To test the efficacy of strict spatial memory without directional persistence, we implemented a Self-Avoiding Walk with a memory capacity of M=1 past spatial coordinates. Step lengths were drawn from an exponential distribution (scale = 1.0), and headings from a uniform distribution (U(0,2π)). To enforce avoidance, any proposed coordinate that fell within a physical Euclidean distance of 0.5 cm from the preceding two nodes was strictly rejected (up to a computational limit of 50 resampling attempts), forcing the agent into unexplored spatial territory.
– Correlated Random Walk (CRW): Representing a standard persistent search without spatial memory, the CRW agent executed continuous 2D angular trajectories. Step lengths were drawn from an exponential distribution (scale = 1.0). The heading angle for each subsequent step was heavily correlated with the previous heading via a bounded Cauchy-like transformation (centered near zero, preventing sharp reversals). The resulting vector was decomposed into its longitudinal (Δz) and transverse (Δθ) components for surface mapping.
– Uncorrelated Random Walk (Brownian Motion): Serving as the null model for spatial memory and momentum, this Brownian-like agent drew step lengths from an identical exponential distribution (scale = 1.0), but its heading angle at each step was drawn entirely independently from a uniform distribution (U(0,2π)). This resulted in highly localized, tortuous trajectories with zero directional persistence.
– Lévy Walk (Scale-Free Search): Representing an optimized search strategy for sparse, unknown targets, the Lévy agent selected step lengths from a heavy-tailed Pareto distribution (shape parameter α=2.0), allowing for clustered local searches interspersed with massive, long-range ballistic relocations. Heading angles were drawn from a uniform distribution (U(0,2π)).

For each of the five search strategies, independent Monte Carlo batches (n=10,000 simulations per strategy) were executed. During each step, the agent’s absolute spatial coordinates were tracked across the curved surface manifold. If the agent’s spatial boundary intersected the boundary of the target hole at any point during its trajectory, the trial was immediately terminated and scored as a success. This procedure was iteratively repeated across an array of discrete target diameters (D∈{0.3,0.5,0.7,1.0,1.5,2.0} cm). The terminal contact probability for each strategy was plotted as a continuous detection curve against hole diameter, enabling a direct benchmark of the agouti’s specialized Markov heuristic against universally recognized theoretical optima (Lévy, CRW, SAW) and random baselines (Brownian). The true empirical success rates of the actual agoutis were overlaid on the resultant curves to validate the predictive power of the generative models.

To evaluate the relative efficiency of the different sensory sampling, we modeled the probability of successful hole detection using a Generalized Linear Model (GLM) with a binomial error distribution and a logit link function (logistic regression). The binomial GLM was structured to estimate the log-odds of successful detection as a function of the underlying navigational strategy and the parametric hole size. The generalized regression equation was defined as:

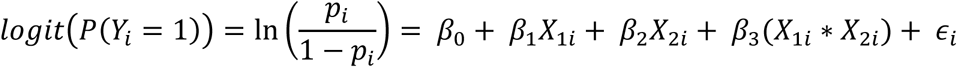

Where:

- *p_i_* is the probability of successful hole detection for simulation iteration.
- *β*_0_ represents the global intercept (the baseline log-odds of detection for the reference group, established as the URW strategy on a True Peanut manifold).
- *X*_1*i*_ is a categorical predictor matrix representing the specific Navigational Strategy employed (Data-Driven, CRW, Lévy).
- *X*_2*i*_ is a continuous predictor representing the parametric Hole Size (ranging from 0.3 cm to 2.0 cm).
- (*X*_1*i*_ ∗ *X*_2*i*_) represents the interaction term between the search algorithm and hole size, testing whether the relative advantage of a specific walk strategy scales non-linearly with the target size.
- *ε_i_* represents the residual error.

The models were fitted using maximum likelihood estimation (MLE). To account for the potential violation of independence across simulations run on identical geometric meshes, we utilized Huber-White cluster-robust standard errors to prevent the artificial inflation of Type I error rates. Model selection and goodness-of-fit were evaluated using the Akaike Information Criterion (AIC) and McFadden’s pseudo-pairwise post-hoc contrasts between the Data-Driven strategy and the null theoretical walks (URW, CRW) were executed utilizing Wald Chi-Square tests on the estimated marginal means (EMMs). To control for the family-wise error rate across multiple strategic comparisons, all resulting *p*-values were adjusted using the Benjamini-Hochberg False Discovery Rate (FDR) correction. An adjusted alpha threshold of 0.05 was utilized to denote statistical significance for the competitive advantage of the empirically derived agouti search algorithm.

##### d. DDM

To investigate how tactile sensory sampling on the surface of an object drives the ultimate binary decision to either “cache” or “eat”, we developed a spatially integrated Drift Diffusion Model (DDM). Unlike standard DDMs where evidence accumulates steadily over time, our architecture explicitly couples the internal decision variable to a physically simulated 3D spatial search. This allows the internal decision bounds to be modulated dynamically by discrete, spatially-dependent sensory events (i.e., encountering the structural hole).

We simulated the physical trajectory of the animal’s teeth across the object’s surface. The model utilizes continuous geodesic random walks over a mathematical 3D representation of the peanut. At each step t (representing a single bite), the spatial coordinates of the virtual teeth are evaluated for intersection with the physical boundaries of the experimentally manipulated hole. If an intersection occurs, a “True Contact” is registered. To account for sensory noise and biological imperfection, the architecture includes a false-positive sensory rate, ɛ, that is fitted but forced to be below 10% to regularize the fitting. At any given step, regardless of the true physical location of the teeth, there is a probability ɛ that the animal incorrectly perceives a structural flaw. Thus, the binary contact sequence C(t) experienced by the internal decision maker is the logical union of true spatial overlaps and stochastically generated false positives. The internal decision state, x(t), is initialized at x(0) = 0. At each step, x(t) is updated according to three forces: an intrinsic drift, sensory-evoked evidence jumps, and Gaussian noise. The recursive update equation is defined as:

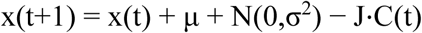

Where:

– μ (Drift Rate): Represents the intrinsic, baseline bias of the animal to cache the seed, independent of any structural flaws. A positive μ constantly pushes the animal towards the caching decision threshold at every bite.
– σ (Diffusion Noise): Represents the inherent stochasticity in the nervous system’s evidence integration. To prevent parameter unidentifiability, noise was fixed proportionally to the drift rate (σ=1.2μ).
– J (Sensory Jump): The massive penalty applied to the decision variable if a sensory contact C(t)=1 is perceived. This negative jump instantly drives the animal away from the caching threshold and towards the consumption threshold.
– B_c_ and −B_e_ (Decision Bounds): The trial terminates when x(t)≥B_c_ (triggering the “Cache” motor program) or when x(t)≤−B_e_ (triggering the “Eat” motor program).

If the animal reaches a biological upper limit of bites (MAX_BITES=100) without crossing a boundary, the decision defaults to the boundary closest to the final state x(t_max_).

The five free parameters of the model (Θ={μ,B_c_,B_e_,J,ɛ}) were fit to the empirical data simultaneously across all experimental hole sizes using the Nelder-Mead simplex algorithm. To ensure the model accurately captured both the final choices and the temporal dynamics of the search, we employed a composite loss function L_total_ that penalized both choice probability errors and step-count distributional divergence:

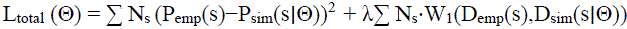

Where s iterates over the experimental hole sizes, N_s_ is the empirical sample size for that condition, P denotes the probability of caching, and W_1_ denotes the 1st-Wasserstein distance (Earth Mover’s Distance) between the empirical distribution of bites-to-decision (D_emp_) and the simulated distribution of steps-to-bound (D_sim_). The scaling factor λ was dynamically initialized during the first iteration to balance the gradients of the sum-of-squared errors (SSE) and the Wasserstein metrics.

Model performance was quantitatively evaluated using binomial Log-Likelihoods.

#### 5. Data analysis and statistics

##### a. Probabilities of Scatter-hoarding

The probabilities of scatter-hoarding that are shown on yellow violin plots with black scatters correspond to distributions of individuals. For each individual the probability of scatter-hoarding a nut of a given condition is computed and this probability corresponds to the black scatter. Once this procedure has been run on all individuals, violin plots are computed as the distribution of scatters, in other words as the distribution of scatter-hoarding probabilities of each individuals.

##### b. Preservability regressions

To assess the structural and biological degradation of the food cache over time, a series of longitudinal putrefaction experiments were conducted on peanuts under three distinct physical conditions mirroring the natural states of cached seeds: structurally intact (control), minimally breached (hole), and fully compromised (crushed). The degradation was quantified across three independent biological and physical metrics: structural resistance (turgidity), fungal colonization (mold proportion), and relative mass loss. For all three metrics, temporal degradation rates were modeled using Ordinary Least Squares (OLS) linear regression. The independent variable in all models was the time elapsed since experimental initiation (measured in days, referred to interchangeably as retrieval day, delay, or recovery time). The slope (β1) of each regression line was extracted to serve as the definitive quantitative measure of the degradation rate for each experimental condition.

The physical integrity of the seeds was measured over time via a structural resistance assay. Resistance values were normalized against baseline measurements taken at Day 0 to account for initial variance in seed hardness. A linear regression model was fitted to the normalized resistance values as a function of the retrieval day. The temporal degradation of structural turgidity was plotted continuously across the experimental window for all three conditions (intact, hole, crushed).

To quantify biological putrefaction and fungal growth, standardized imaging was performed on the retrieved seeds at regular intervals. The extent of fungal colonization was computationally extracted via colorimetric thresholding, specifically isolating the “proportion of red pixels” across the exposed surface area of the seed (as red/brown pixel densities reliably correlated with the specific mold morphologies affecting the caches).

A linear regression model was fitted to predict the proportion of red pixels as a continuous function of the delay (in days). The regression provided the slope (β1, representing the rate of fungal spread), the y-intercept (β0), the coefficient of determination (R^2^), and the statistical significance (p-value) of the growth trend for each of the three structural conditions.

To assess the rate of desiccation and biological consumption of the cache, the absolute mass of the seeds was recorded at standardized recovery times. Because absolute baseline mass varies naturally between seeds, the data was transformed into a proportional weight loss metric relative to the original Day 0 mass.

Similar to the fungal colonization assay, an OLS linear regression (scipy.stats.linregress) was applied to model proportional weight loss as a function of recovery time (days). The regression outputted the rate of mass loss (slope), the R^2^ fit, and the two-sided p-value for the structural conditions. To directly benchmark the differential vulnerability of the three caching conditions, the degradation rates (the slopes extracted from the linear regressions of the mold and mass loss assays) were aggregated and plotted as comparative bar charts. Differences in the magnitudes of these slopes directly illustrate how structural compromises (holes and crushing) accelerate the biological and physical decay of the cached food resource compared to intact controls.

##### c. GLMM

To evaluate the influence of specific behavioral motifs on the target outcome while controlling for individual differences and task conditions, we employed a Generalized Linear Mixed-Effects Model (GLMM).

The relationship between the retained behavioral predictors (independent variables) and the outcome (dependent variable) was modeled using the MixedLM class within statsmodels. We chose a mixed-effects architecture to account for the hierarchical nature of our repeated-measures data, allowing us to estimate population-level trends (fixed effects) while controlling for idiosyncratic animal behaviors (random effects). The model formula was constructed dynamically to include all sanitized behavioral features as fixed main effects. To account for baseline individual differences, the subject identifier (animal_id) was specified as the grouping variable, fitting a random intercept for each animal. Furthermore, to account for condition-specific individual variance, we introduced a random slope for the peanut type (i.e. hole size). This advanced formulation explicitly models the assumption that while the population has a general response to different peanut types, individual animals may exhibit unique, animal-specific sensitivities to the peanut_type condition. Following model convergence, the fixed effect coefficients (estimates), standard errors, and corresponding p-values were extracted to determine the magnitude, directionality, and significance of each behavior’s contribution to the outcome.

